# Divergence of Melanopsin’s Intrinsic Properties and Downstream Actions

**DOI:** 10.64898/2026.09.09.750349

**Authors:** Nguyen-Minh Viet, Franklin S. Caval-Holme, Elliott S. Milner, Andrew S. Johnson, Wendy W. Liu, Alan J. Emanuel, Michael Tri H. Do

## Abstract

Opsins are light-activated receptors. A basic question is how the molecular properties of an opsin shape its effects on the organism. In mammals, melanopsin regulates perception, cognition, physiology, and mood. Because these actions are vital, an international standard has been formulated to express light in terms of melanopsin stimulation. Short wavelengths are absorbed best by melanopsin and should drive its effects most strongly. We report that melanopsin’s cellular and behavioral effects show little or even opposite wavelength sensitivity in some regimes. This divergence stems from melanopsin’s persistent activity, which generates adaptation. Wavelengths absorbed strongly by melanopsin produce high persistent activity and adaptation, giving net desensitization, whereas wavelengths absorbed weakly can produce sensitization instead. We provide a quantitative model that captures these transformations. This work indicates how the downstream actions of a molecule may diverge starkly from its intrinsic properties.

## INTRODUCTION

In the mammalian eye, melanopsin triggers electrical signals that influence perception and are especially critical for driving “non-image” visual functions, such as regulation of the pupil, circadian rhythms, sleep, and mood^1-3^. Like other photopigments, melanopsin absorbs various wavelengths of light to different degrees^4-9^. The axiom is that a photopigment confers its absorption spectrum on downstream processes. This is foundational for understanding light’s biological effects, designing research, and developing artificial lighting. Indeed, melanopsin-driven behaviors have been reported to mirror melanopsin’s absorption spectrum^10-12^. These pioneering studies were conducted when melanopsin was understood to have one ground state and thus one activation spectrum.

In mammals, melanopsin is now recognized to have three highly stable and photoconvertible states^6-9^:

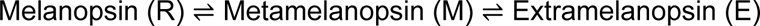

The ground state (“R”), signaling state (“M”), and third state (“E,” electrically silent) have distinct absorption spectra. In studies of purified melanopsin, their spectra peak at 467, 476, and 446 nm, respectively^6^; cellular experiments largely agree^7-9^. Under short-wavelength illumination, the R and E states absorb photons best and therefore depopulate. Because the M state absorbs these wavelengths less effectively, it populates. Thus, at photoequilibrium, the M state dominates. It has high thermodynamic stability; after illumination ceases, it persists and drives cellular activation for minutes^7,8,13^. Conversely, longer wavelengths produce a photoequilibrium in which the M state is depopulated. Different spectra of illumination may therefore produce different levels of melanopsin activation^7,8^. Melanopsin tristability appears unique and highly conserved^8^. It has been recognized for a decade, but its consequences for the organism are largely unexplored.

Prior studies comparing the spectral sensitivity of melanopsin and its downstream processes were designed to minimize the effect of stimulus history. However, stimulus history is a natural influence on sensory responses^14^ and is now understood to have enduring actions on melanopsin signaling. For example, light drives mechanisms of negative regulation—for brevity, "adaptation"—that reduce melanopsin signaling. These include Ca^2+^ influx, phosphorylation, and arrestin binding^15-21^. The availability of melanopsin molecules for photoactivation also changes with light history, as some accumulate in the signaling state and others become insensitive (e.g., by losing their photosensitive chromophores)^7,8,22,23^. How adaptation shapes the expression of melanopsin’s intrinsic molecular properties is unclear^24,25^.

Here, we query the nature of melanopsin signaling in the context of tristability and adaptation. Investigating this topic provides insight into melanopsin, the critical processes it supports, the influence of photopigment properties on the organism, and how lighting may affect health^2,26^. We find that the interaction of tristability and adaptation shapes the manifestation of melanopsin’s effective spectral sensitivity downstream, in behavioral and cellular responses to light. Rather than short and long wavelengths always driving more or less activation, respectively, their effects appear equal or even inverted in some conditions.

## RESULTS

### Effective Spectral Sensitivity of a Melanopsin-Driven Behavior

We examined a behavioral response that depends on melanopsin and operates on timescales that facilitate quantitative study and comparison to melanopsin-driven cellular responses. This is the pupillary light reflex (PLR). It supports vision as light intensifies by sharpening the image and lowering sensitivity^22,27-29^. To focus on melanopsin signaling, we evaluated pupil responses in mice with rod and cone phototransduction disabled (*Gnat1^-/-^; Gnat2^-/-^*)^30,31^. We delivered pulses of light at a frequency that should allow light history to accumulate^7,15,16^. We used brief pulses, approximating impulse stimuli, to focus on system dynamics rather than their convolution with ongoing stimulation^22,32^. We delivered intensities that drove pupil constriction from threshold (the lowest tested intensity corresponds to melanopsin’s photon absorption rate when the sun is 2.5° above the horizon) to beyond the physiological range (melanopsin’s absorption reaches 8 log_10_ photons µm^-2^ s^-1^ in daylight)^33^.

We tested two wavelengths, matching their intensities at the retina by accounting for the eye’s transmission spectrum^33^. The first is short (440 nm) and produces a photoequilibrium of melanopsin states in which the M state dominates; daylight and other kinds of white light produce similar photoequilibria^7^. The second is long (560 nm), which minimizes the M state at photoequilibrium^7^ (**Figure S1**). Exact wavelengths and all details for specific experiments are given in **Table S1**.

We constructed intensity-response (I-R) relations for short- and long-wavelength light (**Figure 1A-D**). After prolonged darkness, all melanopsin holopigment (opsin bound to chromophore) should be in the R state, such that sensitivities to light of these wavelengths should have a ratio of ∼10.7 (440 and 560 nm; **Methods**)^7^. At the foot of pupil constriction’s I-R relation (threshold), we measured a short:long sensitivity ratio of 7.5 (95% confidence interval, CI, of 5.1-9.0; **Figure 1E**). This smaller value is likely due to prior light pulses that activated the pathway but did not evoke detectable constriction^35^. As the I-R relations rise to their saturated maxima, they gradually converge and their response ratio approaches unity (**Figure 1D-E**). In this regard, the spectral sensitivity of melanopsin-driven pupillary constriction naturally diminishes with response saturation.

**Figure 1.**
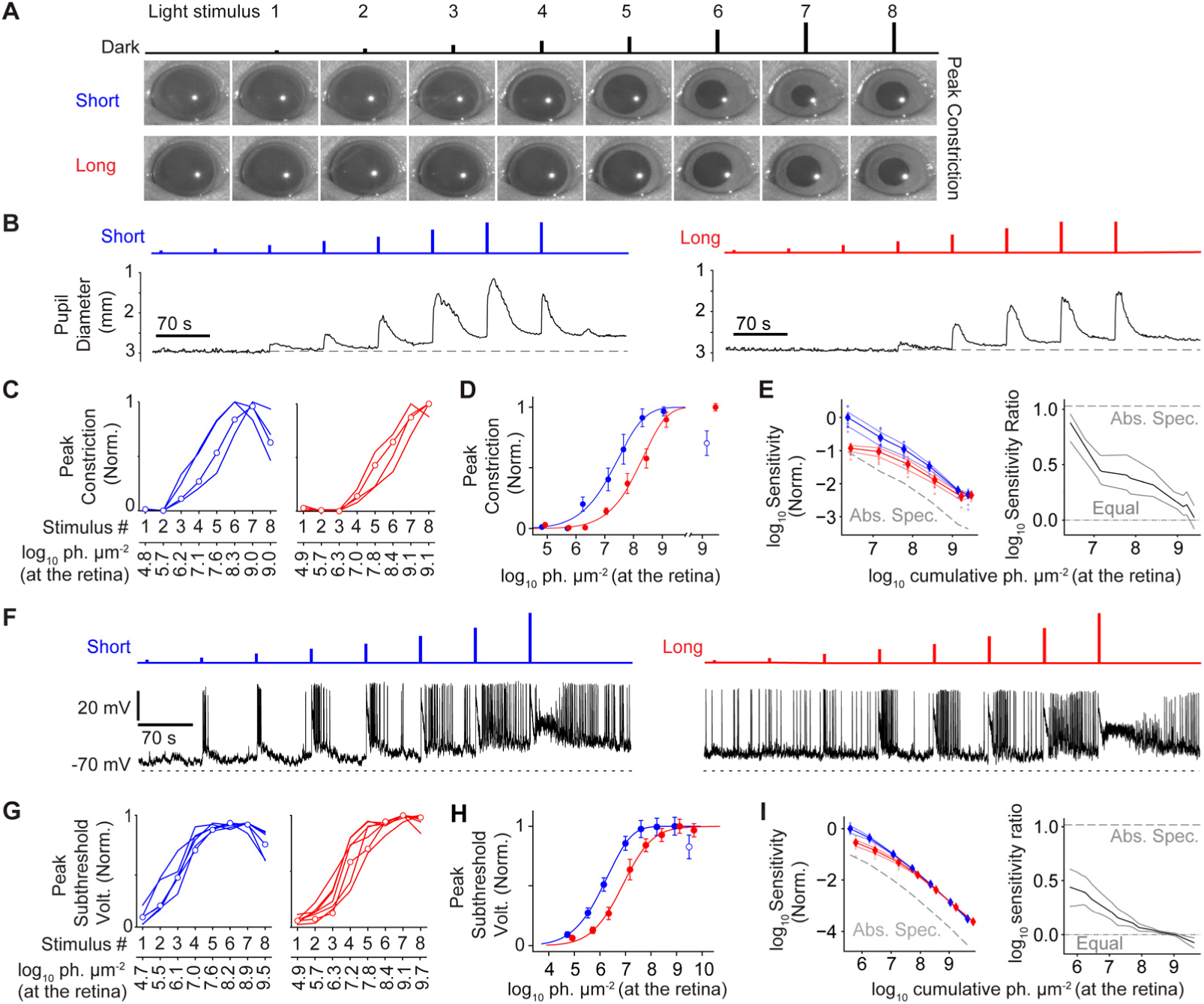
Melanopsin-driven behavior and neuronal activity diverge from melanopsin’s spectral sensitivity. **A.** The pupil in darkness and following pulses of light (stimulus monitor at the top) delivered to the contralateral eye. Here and in subsequent plots, short and long wavelengths are designated by blue and red coloring, respectively. To isolate melanopsin signaling, phototransduction is genetically inactivated in rods and cones. Single trials from one example mouse shown. **B.** Quantification of pupil diameter for single trials from the animal in panel **A**. Upward deflections indicate constriction. The stimulus monitor is at the top. **C.** Intensity-Response (I-R) relations for peak constriction (normalized to maximum). Each curve shows data from one animal (mean of 3 sessions). The short-wavelength stimulus produces a region of pronounced negative slope over higher intensities. Open circles mark the example animal of **A** and **B**. **D.** Mean I-R relations for peak constriction (normalized to maximum mean constriction). The repeated highest intensity is shown on a separate axis to the right. Here and in **H**, open circles indicate regions of negative slope (responses at least 5% smaller than a prior response). Sigmoid fits are to the region of positive slope. Error bars are standard error of the mean. **E.** *Left,* Sensitivity of pupil constriction (mm photons^-1^ μm^2^) under short or long wavelength, normalized to the maximum for the former and plotted against cumulative light history. If long-wavelength sensitivity followed the prediction from melanopsin’s dark-adapted (R state) absorption spectrum (‘Abs. Spec.’), it would fall on the gray dashed line. The prediction for melanopsin’s E state is even lower and is not shown. The first two stimuli are not shown, being consistently below threshold. Individual trials (dots) are averaged (diamonds) and interpolated (lines, with 95% confidence intervals; see **Methods**). *Right,* Short:long sensitivity ratio with confidence intervals. With cumulative light history, the spectral sensitivity shifts away from the prediction from melanopsin’s absorption spectrum (∼1) to equalization (0). **F.** *Bottom,* Patch-clamp recordings of voltage from two example intrinsically photosensitive retinal ganglion cells (ipRGCs, all of the M1 type in this and subsequent figures; 35 °C). *Top,* Timing of the light pulses. **G.** I-R relations of peak subthreshold voltage (normalized). Note the region of pronounced negative slope for that obtained with short-wavelength light. Open circles mark example cells in **F**. **H.** As in D, but for peak subthreshold voltage. **I.** As in **E**, but for peak subthreshold voltage. For each plot, the response to the first pulse was undetectable and not shown. See **Table S1** for additional methodological details and **Table S2** for statistical results. Antagonists of synaptic transmission were included in all *ex vivo* experiments.

We found an additional drive toward spectral equalization. Melanopsin’s absorption spectrum indicates that short-wavelength pulses of increasing intensity should activate larger fractions of melanopsin to evoke stronger pupil constrictions, up to saturation^10,28,29^. Instead, constriction usually peaked at an intermediate intensity and declined thereafter, producing an unusual region of negative slope at the higher end of the intensity-response (I-R) relation (**Figure 1A-D**). Repeating the highest intensity caused constriction to decline even further, to 0.70 ± 0.21 of the maximum (mean ± SD, 3 sessions for each of 4 animals). This indicates adaptation (i.e., negative regulation) because the same stimulus produces an attenuated response. By contrast, long-wavelength pulses of increasing intensity produced increasing pupillary constriction, and repeating the brightest pulse tended to give the same response (0.97 ± 0.06 of the maximum; p = 2 × 10^-4^, Nested Wilcoxon rank test^34^; **Figure 1A-D**). Indeed, for the last pulse tested, absolute pupil constrictions were similar between wavelengths (0.99 ± 0.31 and 1.13 ± 0.22 mm for short and long, respectively; p = 0.27). Thus, when comparing across these lighting conditions, pupil constriction depends far less on wavelength than expected from melanopsin’s absorption spectra.

### Effective Spectral Sensitivity of Melanopsin-Driven Voltage Changes

To investigate where the effective spectral divergence between melanopsin and pupillary constriction originates, we considered that M1 intrinsically photosensitive retinal ganglion cells are key mediators of this process (hereafter, “ipRGCs”)^1,36-38^. These neurons sense light using melanopsin and by receiving synaptic input driven by the classic rod and cone photoreceptors^1^. We examined responses of ipRGCs in the *ex vivo* mouse retina near body temperature. We preserved and isolated their melanopsin-driven responses by using perforated-patch electrophysiological recording and by providing antagonists of synaptic transmission, respectively^7,8,39^. We focused on the subthreshold membrane voltage because of its relatively straightforward dependence on melanopsin phototransduction, as compared to spikes. For comparison with pupillary measurements, we delivered the same stimulus to the retina, accounting for the eye’s transmission spectrum^33^, except for use of a brighter last pulse to probe a broader range of intensities.

The subthreshold photovoltage of ipRGCs was more sensitive than pupil constriction, likely reflecting downstream thresholding of ipRGC signals for the latter^35^. Like pupillary constriction, photovoltage was more similar between wavelength conditions than expected from melanopsin’s absorption spectra. Over higher intensities, photovoltage I-R relations exhibited a stronger region of negative slope for short- than long-wavelength light (p = 0.0082; Wilcoxon rank-sum test for unpaired data unless otherwise noted; **Methods**). At the highest intensity tested, photovoltages were similar across these wavelengths (36.8 ± 10.7 and 39.2 ± 6.0 mV, p = 0.58). Turning to the short:long sensitivity ratio, it was 2.74 (95% CI: 1.84-3.98) at threshold, consistent with photovoltages appearing after some light history. With additional light history, the ratio fell gradually to 0.85 (95% CI: 0.77-0.93; 6 and 7 cells; **Figure 1F-I**). Therefore, from threshold to saturation of ipRGC photovoltage, the difference between short- and long-wavelength conditions progressively diminishes. It then inverts, which is unexpected from melanopsin’s absorption spectra.

### Effective Spectral Sensitivity of Melanopsin Phototransduction

To examine melanopsin phototransduction directly, we clamped the membrane voltage to measure current through the cascade’s TrpC6/TrpC7 cation channels^8,29,40^ and to suppress voltage-sensitive channels. We tested both wavelengths in each cell, examining all four combinations of pulse sets across cells (long-long, long-short, short-short, and short-long; **Figure 2A-C** and **Figure S2**). Doing so shows whether wavelength-specific response properties are robust to light history, and controls for cellular heterogeneity^41,42^. These extended experiments require exceptional stability, so we worked at room temperature, increased inter-stimulus intervals to accommodate the slower kinetics at this temperature^22,41^, and focused on the upper end of the intensity range.

**Figure 2.**
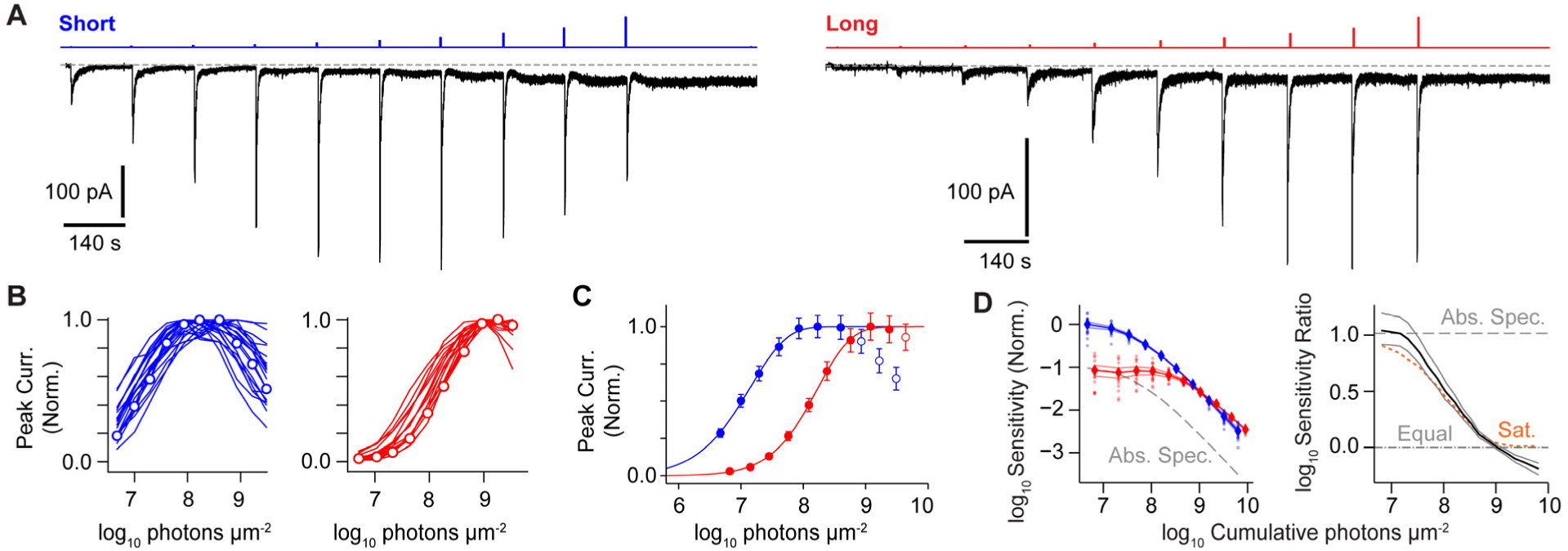
Melanopsin phototransduction diverges from melanopsin’s spectral sensitivity. **A.** Example ipRGCs in voltage clamp (-80 mV) and given intensifying light pulses of either short (*left*; blue) or long wavelength (*right*; red). Dotted lines (gray) mark the pre-stimulus baseline. **B.** Intensity-response (I-R) relations for peak transient current for short and long-wavelength pulse sets. Open circles mark example cells of **A**. **C.** Mean I-R relations for peak current, normalized within each wavelength to the maximum mean response across cells. Open circles mark regions of negative slope (responses at least 5% smaller than a prior response). Sigmoid fits are to the region of positive slope. Error bars are standard error of the mean. **D.** *Left,* Sensitivity (pA photons^-1^ μm^2^, normalized) to the set of short- and long-wavelength pulses plotted against cumulative light history. Measurements from individual cells (points) are averaged (diamonds) and interpolated (lines flanked by 95% confidence intervals; see **Methods**). If long-wavelength sensitivity followed the prediction from melanopsin’s dark-adapted (R state) absorption spectrum (‘Abs. Spec.’), it would fall on the gray dashed line. *Right,* Short:long sensitivity ratio (thick line flanked by 95% confidence intervals). Spectral sensitivities shift from the log ratio predicted by the absorption spectrum (∼1) to equalization (0) and then inversion (< 0). Response saturation alone (‘Sat.’; orange dashed line; see **Methods**) produces equalization, but not inversion. In **A**, 10-mV hyperpolarizing steps (not shown) were given periodically to probe recording parameters and the resulting capacitance transients are often visible. See **Table S1** for additional methodological details and **Table S2** for statistical results. Antagonists of synaptic transmission were included in all experiments.

Overall, melanopsin photocurrent resembled photovoltage and pupil size. Analyzing the first relations for all cells (set 1, initial condition of dark adaptation), we found that negative slope over high intensities was stronger for short- than long-wavelength pulses (p = 6.9 × 10^-6^). At the highest intensity tested, photocurrents were indistinguishable across wavelengths (158 ± 56 and 160 ± 82 pA, p = 0.86; 20 and 15 cells for short and long wavelengths, respectively). The short:long sensitivity ratio began at 10.99 (95% CI: 8.18-15.17), near expectation from melanopsin’s absorption spectrum (**Methods**). With cumulative light history, the ratio fell to a minimum of 0.65 (95% CI: 0.57-0.71; **Figure 2D**). This is the monotonic progression toward spectral equalization, followed by the unexpected inversion where the long-wavelength condition ends with higher effective sensitivity.

Comparing two sets of I-R relations within cells provides a similar view. For instance, negative slope was pronounced for short-wavelength light whether it was used for the first set, second set, or both sets of pulses (**Figure 3A-B** and **Figure S2**). Likewise, negative slope was milder or absent for long-wavelength light whenever it was used (**Figure S2**). Therefore, the divergence of effective spectral sensitivity between the melanopsin molecule and its phototransduction cascade is robust to light history and cellular heterogeneity.

**Figure 3.**
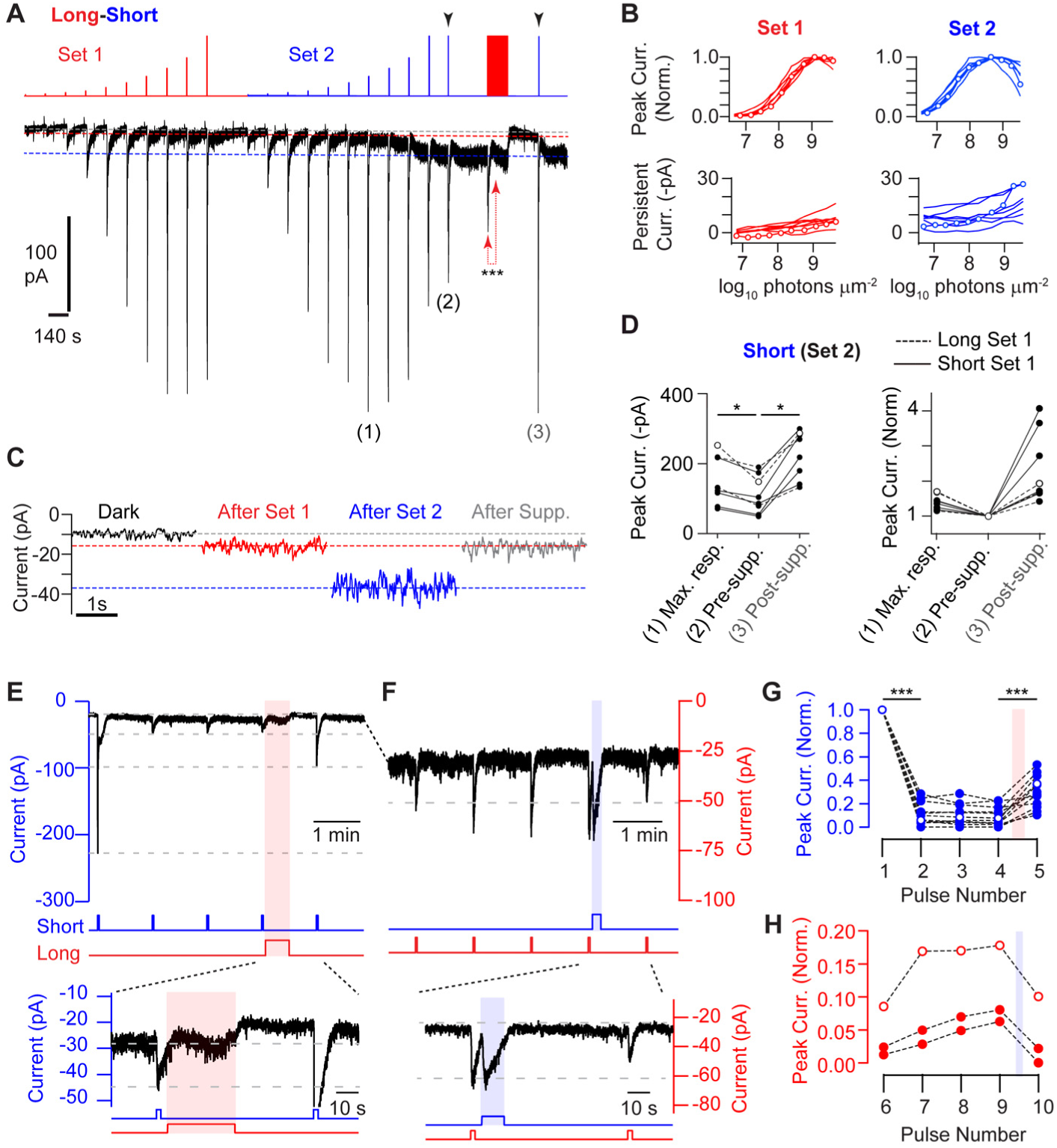
Dependence of melanopsin phototransduction on persistent activity and adaptation. **A.** Example ipRGC given two consecutive sets of intensifying light pulses of long or short wavelength. The cell received an additional pair of high-intensity, short-wavelength pulses (black arrowheads) with an intervening long-wavelength pulse to suppress persistent activity. During this long-wavelength suppression pulse, the current was initially large and then declined, evincing adaptation (connected red arrows; observed in 15 additional cells). The transient response amplitude was largest at an intermediate intensity (1), smaller at the repeated highest intensity (2), and recovered by the long-wavelength period (3). Dotted lines mark the pre-stimulus baseline (gray), magnitude of the largest persistent activity evoked by long wavelength (red), and that evoked by short wavelength (blue). **B.** Intensity-response relations for peak transient (*top*) and persistent current (*bottom*) for the stimulus in **A**. Open circles mark the example cell. **C.** Excerpts of persistent currents from panel **A**, measured in a 3-s window prior to the next light pulse. “After Supp.” is after the long-wavelength pulse that suppressed persistent activity. **D.** *Left,* Peak transient photocurrents from short-wavelength, second pulse sets. Shown are the maximum response to any pulse before long-wavelength suppression (1) and the response to the pulse preceding suppression (2) or after suppression (3). Solid and dashed lines indicate cells for which the first pulse set was short- or long-wavelength, respectively. Open circles indicate the example cell in **A**. *Right,* The same transient photocurrents normalized within-cell to the pre-suppression transient response. **E.** Repeated, saturating pulses of short-wavelength light with an intervening period of long-wavelength light to suppress persistent activity. *Inset,* Expanded view of the two short-wavelength pulses bracketing the long-wavelength pulse. Dashed lines are for comparison of response magnitudes. **F.** Continuation of the experiment for the cell in **E** (<1 min delay). Persistent activity is ongoing. Long-wavelength pulses, calibrated to be sub-saturating, are delivered with an intervening short-wavelength pulse. *Inset,* Expanded view of the two long-wavelength pulses bracketing the short-wavelength pulse. Dashed lines are for comparison of response magnitudes. **G.** Response magnitudes, normalized to the first, for all 12 cells tested with short-wavelength pulses (example shown as open symbols). **H.** Same as **G** but for long-wavelength pulses. Red and blue shadings indicate timings of intervening long- and short-wavelength pulses, respectively. In **A**, 10-mV hyperpolarizing steps (not shown) were given periodically to probe recording parameters and the resulting capacitance transients are often visible. See **Table S1** for additional methodological details and **Table S2** for statistical results (* p < 0.05, ** p < 0.01, *** p < 0.001). Antagonists of synaptic transmission were included in all experiments.

### Persistent Activity Produces Adaptation

Examination of melanopsin photocurrent suggests how sensitivities under short- and long-wavelength conditions come to appear similar. Short-wavelength pulses of increasing intensity evoked cumulative increases in persistent photocurrent—that which long outlasts the stimulus (also see **Figure S3A**)^7,8^. At higher intensities, this increase was accompanied by a drop in the transient photocurrent, which is reflected in the I-R relation’s region of negative slope (**Figure 2A-C**). By contrast, long-wavelength pulses caused little accumulation of persistent photocurrent and the transient photocurrent magnitude increased to a plateau (**Figure 3A-D**). These observations suggest that persistent activity shapes the I-R relation.

To test this idea, following the second set of short-wavelength pulses, we delivered a period of long-wavelength light (100-140 s) to depopulate the M state and thus suppress persistent current (**Figures 3A**, **3C**, **S1**, and **S3A**). This suppression increased the transient current evoked by a subsequent short-wavelength pulse (potentiation of 236 ± 95%, n = 8 cells; **Figure 3D**; also see **Figure S2D**). Suppression itself drove adaptation, as expected^15,16,41^ and as indicated by the decay of photocurrent during the period of long-wavelength illumination (**Figure 3A** and **Figure S3A**). Nevertheless, the net effect of suppression is sensitization. These experiments indicate that short-wavelength light produces high persistent activity, which drives adaptation that in turn reduces responsiveness.

In separate experiments, we delivered a sequence of identical, short-wavelength pulses that should each drive a saturated response, reaching a photoequilibrium with high occupancy of melanopsin’s signaling M state (**Figure 3E** and **G**; also see **Figure S1**). The first pulse induced a transient response and persistent activity. The following pulses evoked transient responses, riding atop persistent activity, that were attenuated or undetectable (9 and 3 of 12 cells, respectively). We then delivered a period of long-wavelength light, which suppressed persistent activity (10 of 12 cells). A subsequent short-wavelength pulse produced a larger response; it also restored persistent activity (in those 10 cells).

We then delivered long-wavelength pulses that were each calibrated to partially suppress persistent activity, so that melanopsin fractions should proceed incrementally toward a photoequilibrium where M-state occupancy is low (**Figure 3F** and **H**; also see **Figure S1**). Each decrement of persistent activity was accompanied by a larger transient response to the next pulse (**Figure 3H**). A subsequent short-wavelength pulse restored persistent activity, and the transient response to the next long-wavelength pulse was diminished (3 of 3 cells). Persistent activity appears to drive graded levels of adaptation in ipRGCs (also see **Figure 2A** and **Figure S2**).

### Adaptation Shapes Persistent Activity

Having found evidence that persistent activity produces adaptation, we asked if adaptation acts in turn on persistent activity. We previously observed that Ca^2+^ influx drives adaptation in melanopsin phototransduction, such that its acute removal increases the sensitivity and magnitude of light responses in ipRGCs^16^. Therefore, we removed extracellular Ca^2+^ and asked if persistent activity increased. We continued to use voltage clamp and synaptic antagonists to isolate the effect of this manipulation on phototransduction^16^. Persistent activity doubled in size (increasing from 8.5 ± 4.8 to 16.9 ± 7.2 pA, 9 cells; **Figure 4A, B** and **D**). For an internal control, we measured dim-flash responses in the same experiment. A dim-flash response is in the linear range of phototransduction; its waveform reflects that of a single-photon (single-melanopsin) response and its amplitude, divided by the flash intensity, gives a measure of absolute sensitivity (S_F_, in pA photons^-1^ µm^2^)^22,43^. Ca^2+^ removal did not change the initial trajectory of these responses, indicating little or no dependence of activation on Ca^2+^ influx. However, the response in 0 Ca^2+^ rose further to reach a ∼2-fold larger peak, indicating that Ca^2+^ influx usually curtails activation, as expected from a mediator of adaptation (i.e., driving a reduction of activity; **Figure 4A, C** and **E**). These experiments indicate that adaptation strongly regulates persistent activity (also see **Figure S3**).

**Figure 4.**
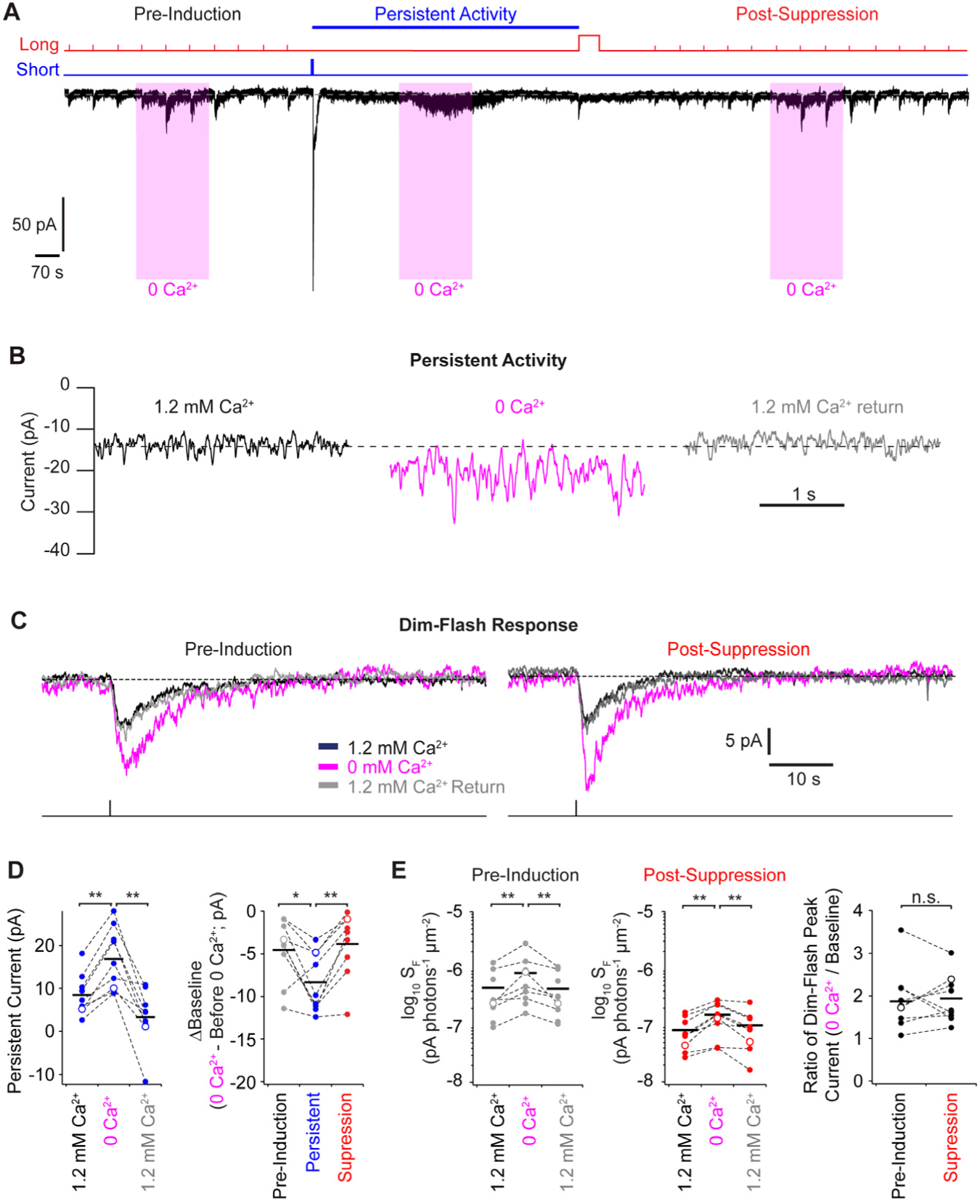
Adaptation shapes persistent activity. **A.** Protocol to evaluate the effect of Ca^2+^ removal on persistent activity (*top*) and its effect on an example cell (*bottom;* same cell in **B** and **C**). A pre-induction period is followed by induction (short-wavelength pulse) and then suppression (long-wavelength pulse) of persistent activity. In each period, extracellular Ca^2+^ is acutely removed and then returned, and the effect on the holding current (which includes any persistent activity) assessed. In the pre-induction and post-suppression periods, dim-flash responses are assessed. 10-mV hyperpolarizing steps (not shown) were given periodically to probe recording parameters and the resulting capacitance transients are sometimes visible. **B.** Effect on the persistent current of removing and replenishing extracellular Ca^2+^. **C.** Effect on the dim-flash response of extracellular Ca^2+^ manipulations in pre-induction (*left*) and post-suppression (*right*) periods. **D.** *Left,* Persistent activity (the change in holding current after the induction pulse) increases in 0 Ca^2+^. *Right,* 0 Ca^2+^ increases the holding current before induction and after suppression of persistent activity, but this increase is greater during persistent activity, indicating an effect on persistent activity itself. **E.** In the same sample as **D**, dim-flash sensitivity (S_F_) increases in 0 Ca^2+^ whether in the baseline or post-suppression periods (*left* and *middle*, respectively). The effect of 0 Ca^2+^ is similar between these periods (*right*). See **Table S1** for additional methodological details and **Table S2** for statistical results (* p < 0.05, ** p < 0.01, *** p < 0.001). Antagonists of synaptic transmission were included in all experiments.

### Transformations of Melanopsin Signaling by the Interaction of Persistent Activity and Adaptation

Persistent activity appears to produce adaptation, and adaptation appears to regulate persistent activity. We sought to predict how this interaction shapes the effective spectral sensitivity of melanopsin signaling, iterating across three models of increasing complexity and accuracy. We began with our existing computational model of melanopsin’s three photoconvertible states (R, M, and E; **Figure 5A** and **Figure S1**)^6,7^. This model produces persistent activity, in the form of M state occupancy that remains after illumination ceases. It also incorporates some adaptation. For example, as molecules are activated to the M state, fewer are available for activation. Moreover, long-wavelength light increases the fraction of E, which reduces responsiveness to wavelengths that this state absorbs more poorly than the R state. This model captures the known photoconversions of melanopsin.

**Figure 5.**
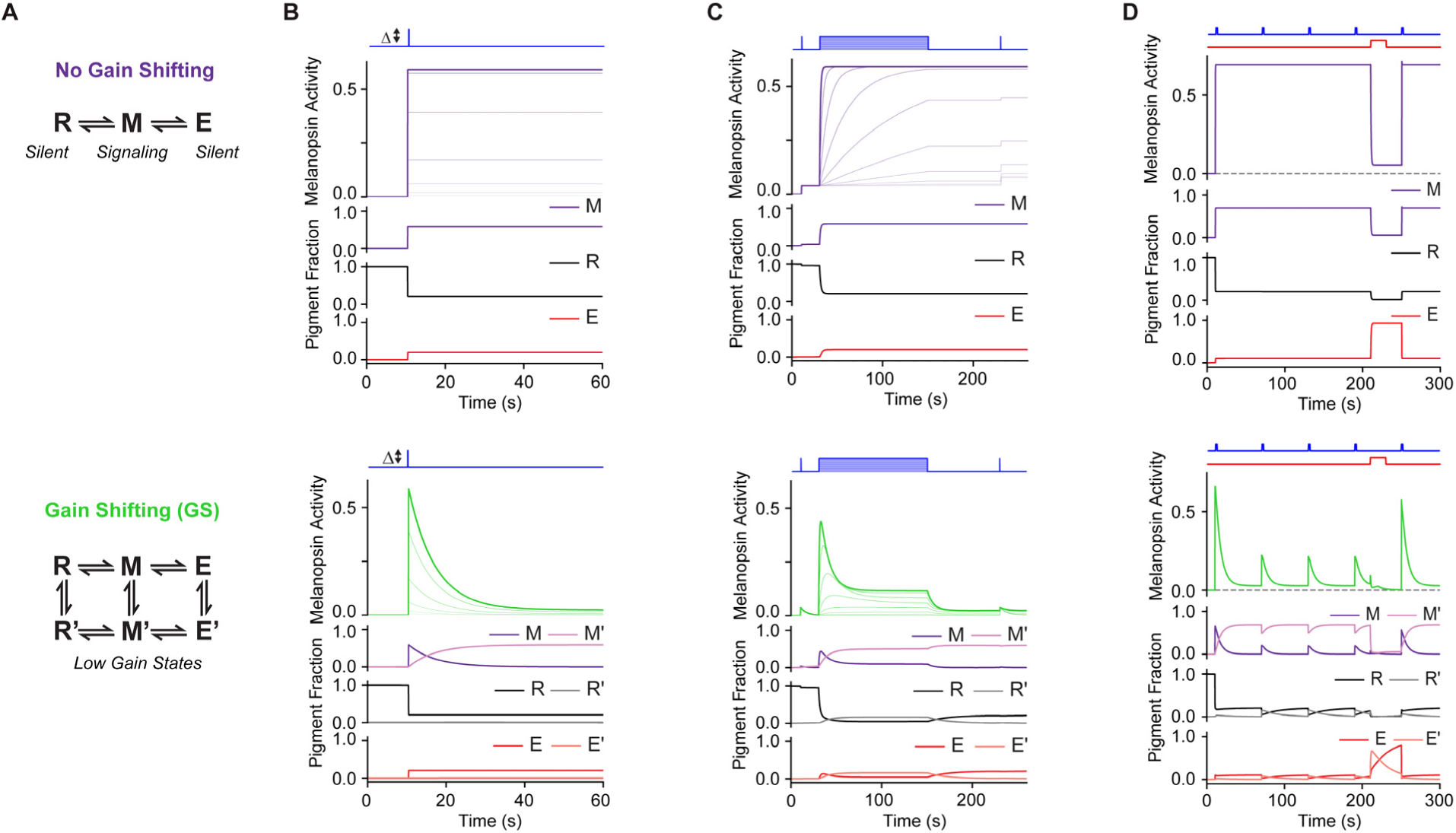
Gain shifting is required for steady irradiance encoding. **A.** Schematics of melanopsin models without (*top row*) and with (*bottom row*) adaptation implemented as shifts to low-gain states (R’, M’, and E’). Horizontal transitions are light-dependent, while vertical transitions are light-independent. “Melanopsin activity” is the fraction of molecules in the M state (*f*^*M*^) for the no gain-shifting (no GS) model. For the GS model, it is *f*^*M*^+ 0.038*f*^*M*′^ (see **Methods**). **B.** Simulated melanopsin activity for a set of flashes varying in intensity (100 ms, 7-10 log_10_ photons µm^-2^ in 0.5-log_10_ increments). The addition of gain shifting (*bottom*) allows for the activity to decay. **C.** Without gain shifting, steps (3-8 log_10_ photons µm^-2^ s^-1^ in 0.5-log_10_ increments) do not produce steady activity that grades with light intensity. Flashes (200 ms, 7.5 log_10_ photons µm^-2^) are provided for reference. **D.** The protocol used in Figure 3E (see **Table S1** for stimulus details). When persistent activity is saturated, subsequent pulses do not generate additional changes in melanopsin activity (*top*) unless gain shifting is implemented (*bottom*).

We found four model behaviors that indicate a need for additional adaptation. First, flashes produce stepwise increases in the M fraction and thus phototransduction (**Figure 5B**). Measured flash responses differ, having a large transient component^22^ (**Figure 4**). Second, different intensities of sustained light produce the same steady-state level of phototransduction, which is set by the photoequilibrium fraction of M (**Figure 5C** and **Figure S1**). By contrast, measured step responses settle at steady levels that grade with light intensity^16,41^. Third, activity during illumination is the same magnitude as that persisting in subsequent darkness (**Figure 5B-C**). Cellular measurements show that persistent activity is far smaller^7^ (**Figure 2A-C** and **Figures S2** and **S3**). Fourth, pulses of light given during saturated persistent activity produce no response (**Figure 5D**). In actuality, “increment” responses can be driven atop saturated persistent activity^7^ (**Figure 3E**). These observations underscore the divergence between melanopsin’s photoconversions, which the model simulates, and their cellular effects.

We therefore added to the model a tier of adapted states (R’, M’, and E’; **Figure 5A**), which may be considered analogous to those produced by forms of adaptation like receptor phosphorylation and arrestin binding^17-21,44^. Molecules in the M state drive phototransduction; those in the M’ state do so with low gain (little activation per molecule); and those in the remaining states do not (no activation per molecule). The magnitude of phototransduction increases with the fraction of molecules in M and M’. This simple, “gain shifting” (GS) model accounts for the four features of melanopsin phototransduction mentioned above (**Figure 5B-D**). First, flash responses are transient because M, once formed, transitions into the low-gain M’ state (**Figure 5B**). Second, during sustained illumination, melanopsin activity reaches steady levels that grade with light intensity (**Figure 5C**). This outcome depends on a dynamic equilibrium of melanopsin states. At low intensities, molecules in the M’ state are photoconverted into R’ and E’ at low rates, yielding little recovery to R and E. Because of this and the low photon absorption rate, photoconversions of R and E to the high-gain M state are sparse. On the other hand, at high intensities, the cycling of molecules from the low-gain M’ state to the high-gain M state (via R’/E’ and R/E) is faster. In other words, during illumination, melanopsin activity is greater in brighter light because the system is operating at higher, continually replenished gain—a process of “gain cycling.” Third, persistent activity is smaller than the preceding activity during illumination. This is because persistent activity arises entirely from the low-gain M’ state, while activity during illumination has gain cycling (**Figure 5C**). Fourth, light given during persistent activity drives gain cycling, producing increment activity (**Figure 5D**). In addition, the GS model mimics recovery of the saturated response following suppression of persistent activity (**Figure 5D**; see **Figure 3** for experimental data). To summarize, simulating principal features of melanopsin phototransduction requires balancing persistent activity with adaptation.

Despite the GS model’s strengths, it misses aspects of the empirical I-R relations (**Figure 6B-E**): modeled persistent activity saturates at relatively modest intensities, negative slope at the top of the relation is mild or absent under short-wavelength illumination, the short:long sensitivity ratio diminishes with light history but does not invert, and response kinetics are oversimplified. Augmenting the GS model reproduces these features (**Figure 6A-H**). Adding loss and recovery of melanopsin (analogous, for example, to pigment bleaching/regeneration^20,22,23,45^) gives appropriately graded levels of persistent activity. Allowing persistent activity to drive adaptation, thus reducing responses to additional illumination, produces negative slope under short- but not long-wavelength light (**Figures 2-3**; see **Figures S2** and **S3** for additional evidence that persistent activity drives adaptation). Incorporating an additional adaptation mechanism, based on total activity, improves response kinetics (**Figure S6**). This augmented GS (aGS) model fits the data well (**Figure 6B-H** and **Figures S4** and **S5**). It is as accurate as any single ipRGC’s measured I-R relation for predicting the population’s I-R relations (**Figure 6E** and **Figure S10**). Accordingly, the aGS model recapitulates the short:long sensitivity ratio and its change with cumulative light history (from 7.9 to 0.57; **Figure 6G-H**). It also recapitulates shifts in spectral sensitivity previously measured in ipRGCs following conditioning with long-wavelength light^7^ (**Figure S7**). The need for complexity in the aGS model provides a rationale for the numerous mechanisms of adaptation observed in melanopsin signaling^17,18,20-23,44,46-49^ (**Figures S5** and **S6**).

**Figure 6.**
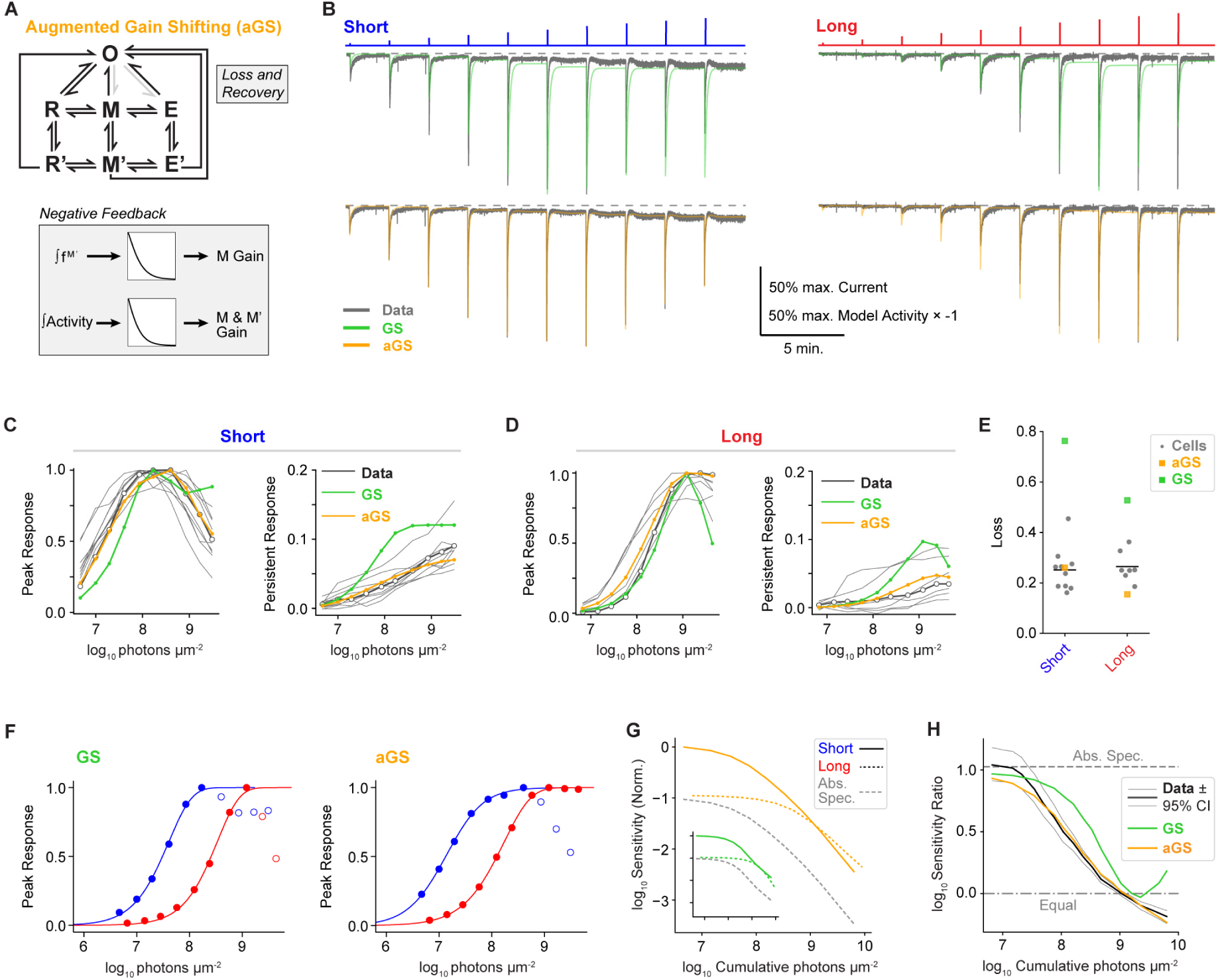
An augmented gain-shifting model predicts spectral divergence. **A.** Schematic of the augmented GS model (aGS). *Left*, All states transition to a non-photoconvertible state (O) that has minuscule signaling gain and gradually returns to R. Light gray arrows indicate O’s potential return to E and M (disabled here; see **Methods**). *Right*, Two gain controls accumulate through leaky integration of persistent activity, corresponding to the fraction of melanopsin in M’ (∫f^M’^) and the summed activity of all states (∫Activity). A nonlinearity governs each control’s reduction of melanopsin signaling gain. See **Methods** for further details. **B.** Model responses (GS: green; aGS: orange) inverted for comparison to example ipRGC photocurrents (gray; data are from Figure 2). Short- and long-wavelength pulse series are on the left and right, respectively, with light monitors at the top. **C.** Short-wavelength intensity-response (I-R) relations for the GS (green) and aGS (orange) models, compared with those of ipRGCs (gray lines). See **Figure S5** for other stimuli. **D.** Same as **C**, for long-wavelength pulse series. **E.** Loss (mean squared error between predicted and cellular I-R relations) with predictions from the GS model (green squares), aGS model (orange squares), or individual ipRGCs (gray dots). Black bars indicate the mean loss for individual cellular predictions. **F.** I-R relations for short- (blue) and long-wavelength (red) pulse series predicted by the GS (*left*) and aGS (right) models. Open circles indicate regions of negative slope (responses at least 5% smaller than a prior response). Sigmoid fits are to the region of positive slope. **G.** Simulated sensitivity (normalized to sensitivity at the first short-wavelength pulse) of GS (*inset*; green lines) and aGS (orange lines) responses to the short- and long-wavelength pulses (solid and dashed lines). Light history is expressed as a cumulative photon count. Predictions (gray dashed lines) are the short-wavelength sensitivity of GS and aGS models divided by the empirical short:long sensitivity ratio for melanopsin’s dark-adapted absorption spectrum (R state; 10.7; see **Methods**). **H.** Short:long sensitivity ratios predicted by the GS model (green line) and aGS model (orange line) compared to the measured ratios from voltage clamp experiments (heavy black line, from Figure 2). Thin lines are 95% confidence intervals. With cumulative light history, relative spectral sensitivity shifts away from that predicted by melanopsin’s absorption spectrum (‘Abs. Spec.’; ∼1) to equalization (0) and then inversion (< 0).

### Transformations of Melanopsin Signaling under Naturalistic Illumination

We have used short and long wavelengths to probe the spectral sensitivity of melanopsin and its downstream actions, but common light sources have a broad spectrum. Thus, we tested xenon (Xe) light, which has a spectrum resembling that of daylight^33^. We modified its spectrum to mimic filtering by the eye’s optics^50^, matched its intensity to that of the short-wavelength light used in our other experiments, and delivered it to ipRGCs (**Methods**). The I-R relations were qualitatively similar to those elicited by short-wavelength light, including high persistent activity and a region of prominent negative slope at higher intensities (**Figure S8**). These features were robust, being similar for Xe delivered without ocular filtering, which enriches for short wavelengths^50^ (**Figure S8**). Our aGS model simulates these effects of Xe light accurately (**Figure S9**). These observations indicate that persistent activity and adaptation shape melanopsin signaling under natural conditions.

## DISCUSSION

We find that the spectral sensitivities of cellular and behavioral responses can diverge from the absorption spectrum of the photopigment that drives them. In the case of melanopsin, which drives many vital processes^1,2,26,51,52^, this divergence results from an interaction of persistent activity and adaptation, both of which follow from light history^2,26,51^. The interaction makes melanopsin-driven responses more agnostic to wavelength than melanopsin itself. This favors encoding of the overall light level rather than image details (i.e., spectral composition). Such downstream transformations of photopigment activity are likely to be important considerations in studies of the visual system, lighting design, and optogenetic applications.

Persistent activity reflects melanopsin’s stable activation^7,8^. This stability has advantages; for example, it blurs spatiotemporal contrasts to favor encoding of the overall light intensity^7,8^. It also can be problematic because light of any intensity eventually causes the same number of activated melanopsin molecules to accumulate, which contrasts with the role of ipRGCs in steadily encoding light intensity^8,28,42,53^. Our study indicates that intensity encoding may emerge from the cycling of gain in a dynamic photoequilibrium of melanopsin states. Gain cycling could exist beyond ipRGCs, such as in invertebrate photoreceptors that use photoconvertible pigments and display adaptation^54^. Indeed, persistent activity and adaptation are widespread and arise from molecular, cellular, circuit, and systems mechanisms^14,55,56^. Their interaction may shape biological signaling in diverse contexts.

Additional investigations may deepen the understanding of melanopsin signaling in context. The challenges include obtaining a quantitative understanding of melanopsin phototransduction’s many mechanisms of adaptation and their influences^15-21^; learning how these mechanisms interact with ipRGC properties that are dynamic, exhibit unconventional features like depolarization block, and span timescales from milliseconds to hours^8,42,53^; capturing the variations of environmental illumination that occur across the day, seasons, and latitudes^57^; and defining how animals sample these conditions^58,59^. The experimental observations and computational model provided here are a framework for such efforts. They also support the rational design of lighting to support research and promote health.

## Acknowledgments

We thank Mark Andermann for expertise on *in vivo* experiments; Ofer Mazor and Pavel Gorelik for the design and construction of devices; Marie Burns, Vladimir Kefalov, and King-Wai Yau for mouse lines; and Chinfei Chen, Marla Feller, Paul Gamlin, Samuel Brill-Weil, Andreas Liu, Philippe Morquette, Navid Mousavi, Victoria Amstrup Vold, and Sophia Wienbar for comments on the manuscript. Funding was provided by the National Institutes of Health (EY023648, EY034089, EY025555, EY032731, and EY036071 to MTHD; EY025466 to ESM; EY033639 and EY037374 to FSC-H; HL007901 to AJE; 1U54HD090255 to the Boston Children’s Hospital IDDRC; and P30 EY012196 to Harvard Medical School); the National Science Foundation (GRFP to AJE); and private foundations (Whitehall Foundation, Karl Kirchgessner Foundation, Knights Templar Eye Foundation, March of Dimes, and Alcon Research Institute to MTHD). MTHD is affiliated with the Center for Brain Science (Harvard University), the Division of Sleep Medicine (Brigham and Women’s Hospital, Harvard Medical School), and the Broad Institute of MIT and Harvard. All authors declare no competing interests.

## Author contributions

Conception: AJE, MTHD. Experiment design: N-MV, FC-H, AJE, MTHD. *Ex vivo* experiments: N-MV, AJE, ESM. Pupillometry: FC-H. Computational modeling: FC-H, ASJ, AJE. Analysis and interpretation: N-MV, FC-H, ESM, AJE, MTHD. Preliminary experiments: AJE, WL. Writing: FC-H, AJE, and MTHD with input from all authors.

Materials are available upon request to. Code is available on GitHub (https://github.com/MichaelDo-Lab/melanopsin-model).

## Use of Artificial Intelligence

The authors used ChatGPT and Cursor for aid with literature searches, proofreading, and coding. The authors carefully vetted the outputs of these models and revised them as needed. The authors assume full responsibility for all aspects of this manuscript.

## Supplemental Information

### KEY RESOURCES TABLE

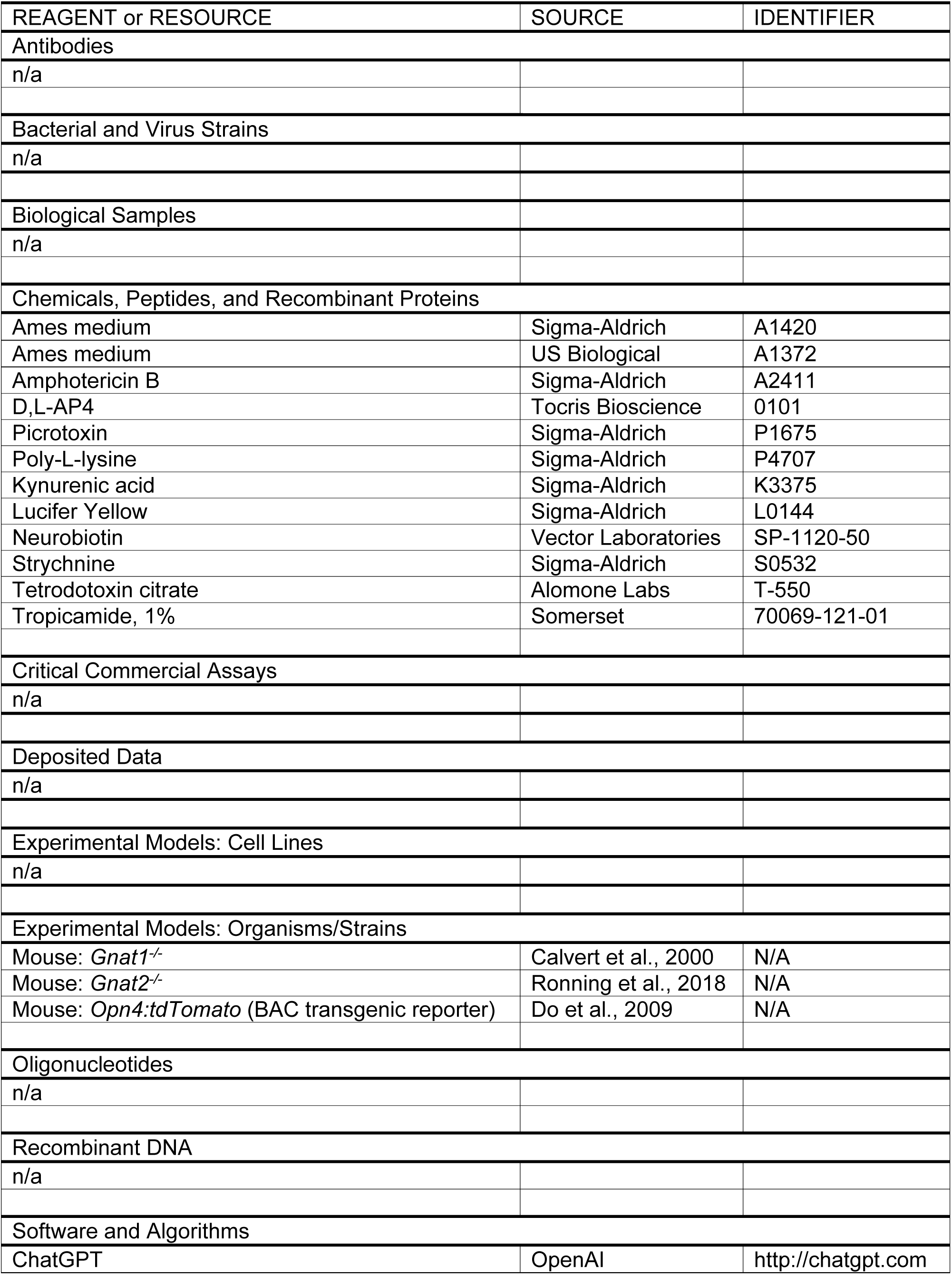
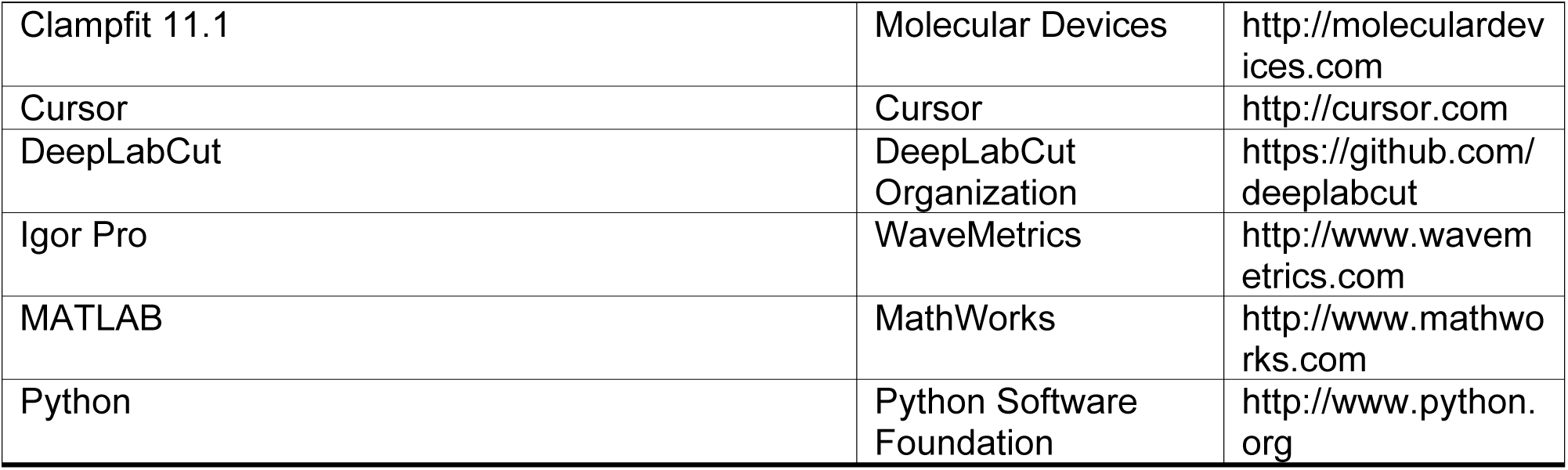

### Methods

#### Animals

All animal procedures were approved by the Institutional Animal Care and Use Committee of Boston Children’s Hospital. For *in vivo* experiments, mice (≥P60) lacked components that rods and cones require for phototransduction (*Gnat1^-/-^* and *Gnat2^-/-^*, respectively)^30,31^. For *ex vivo* experiments, mice (≥P21) were BAC-transgenics in which tdTomato is expressed from the melanopsin gene locus of a bacterial artificial chromosome^22^. M1 ipRGCs are labeled and endogenous melanopsin alleles are not targeted. Animals were maintained on a 12:12 light:dark cycle. Males and females were used. No overt variation with circadian time, sex, or age was observed.

#### Spectral Sensitivity of Melanopsin

The standard for describing the spectral sensitivity of a photopigment is the mathematical template of Govardovskii and colleagues^4^, which is accurate in the majority of cases. When Shichida and colleagues examined purified melanopsin, they found that the R and M state absorption spectra were broader than the Govardovskii template^6^. The action spectrum measured from ipRGCs following dark adaptation, when all melanopsin holopigment should be in the R state, also appeared broader than the Govardovskii template^7,9^. A broadened template, fit to the R state absorption spectrum, also fits the dark ipRGC action spectrum (**Figure S1**).

Here, this unusual spectral sensitivity was tested further using perforated-patch, voltage-clamp recordings from mouse M1 ipRGCs. To promote dark adaptation, ipRGCs were kept in darkness for ≥10 min following fluorescence identification. Previously, this sufficed to make R the only state detectable in ipRGCs by electrophysiology^7^, consistent with biochemical experiments using longer dark adaptation of the animal^60^. As an additional precaution, the temperature was raised to 35 °C in this period. This heat cycle appears to help reset phototransduction; for example, it accelerates the decay of any persistent activity remaining from fluorescence identification (produced by melanopsin’s M state; not shown). Synaptic antagonists were added to isolate melanopsin activity^7,22^, and ipRGCs were stimulated with dim flashes. Dim-flash sensitivities for 440- and 560-nm light had a ratio of 10.6 ± 0.6 (6 cells). This ratio matches that measured previously from ipRGCs without heat cycling (10.7 ± 5.0, 6 cells) ^7^. Thus, the dark-adapted action spectrum of the ipRGC intrinsic response is more consistent with purified melanopsin’s broad absorption spectrum (15.3) than with the narrower Govardovskii template (37.0 with λ_max_ = 467 nm)^4,6^. The ratio of 10.7 is used in the present work.

The spectral sensitivities of the R and M states were fit with the spectral template:

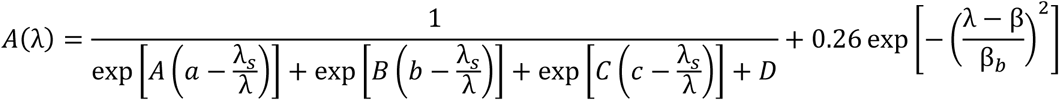

where

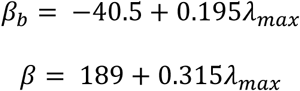

λ_s_ influences template shape but does not determine the exact peaks of the R and M non-Govardovskii spectral templates. The peak spectral sensitivity values (λ_max_) for R, M, and E are 467, 476, and 446 nm, matching those reported for purified melanopsin^6^.

Parameters for the R state were fit to action spectra of dark-adapted ipRGCs^7^:

| **Parameter** | **Value** |
| --- | --- |
| **A** | 44.58 |
| **a** | 0.93 |
| **B** | 8.61 |
| **b** | 0.99 |
| **C** | -11.59 |
| **c** | 1.13 |
| **D** | 0.0 |
| $\lambda_s$ | 494 |
| $\lambda_{max}$ | 467 |

Parameters for the M state were from Shichida’s fit to purified melanopsin^6^:

Parameters for the M state were from Shichida's fit to purified melanopsin<sup>6</sup>:
| **Parameter** | **Value** |
| --- | --- |
| **A** | 44.40 |
| **a** | 0.91 |
| **B** | 11.77 |
| **b** | 0.87 |
| **C** | -14.09 |
| **c** | 1.14 |
| **D** | 0.26 |
| $\lambda_s$ | 483 |
| $\lambda_{max}$ | 476 |

Melanopsin’s E state was fit with a Govardovskii template (with the *a* parameter adjusted based on λ_max_)^4^. There is no published alternative. While modeling intensity-response (I-R) relations (see below), several candidate templates were explored for E but none performed as well as the Govardovskii template (not shown). A broadened template of the form used for R underestimated the 440:480 sensitivity shift following long-wavelength illumination (not shown), whereas a model using the Govardovskii template captured it well (**Figure S7**).

#### Pupillometry

Mice (littermates; one female) were surgically implanted with headplates and individually housed. After a >2-week recovery, mice were acclimated to handling and then to the experimental apparatus (>3 sessions, 20-30 min/session). Prior to each experimental session (in which one I-R series was obtained), mice were dark adapted (>1 hr). All sessions took place 2-8 hrs into the dark phase of the 12:12 hr light:dark cycle. Mice were head-fixed on a passive treadmill fitted with a locomotion encoder. The stimulated (right) eye was dilated with topical application of tropicamide for precise control of retinal illumination. The imaged (left) eye was monitored under infrared illumination (940 nm, 70 nm width at half-maximum, insufficient for retinal activation). Mice received food in their home cages at the end of each session. A pair of points along the iris muscle were tracked by DeepLabCut and the pupil’s diameter measured between them. Summary statistics reported in the main text were calculated across sessions, with a matched number of sessions per animal. Computing these statistics across animals yielded similar measurements. For short-wavelength light, the ratio of constriction at maximum intensity to maximum constriction was 0.7 ± 0.21 (mean ± SD) across sessions vs. 0.7 ± 0.19 across animals.

#### *In Vivo* Optical Stimulation

Light from LEDs (Thorlabs Solis 445C and Solis 565C) was conditioned by 10-nm bandpass filters centered at 440 and 560 nm, respectively. Intensity was controlled by pulse-width modulation (20 Hz; an equivalent intensity of continuous light evoked indistinguishable pupil constriction, not shown) and neutral density filters. Uniform spots of light (8-mm diameter) were projected onto the stimulated eye, overfilling the fully dilated pupil. To estimate the intensity of light at the retina, the intensity at the cornea was multiplied by the ratio of pupillary and retinal surface areas (∼0.43, given a 7.1 mm^2^ fully dilated pupil and a 16.5 mm^2^ retina^61^). Differential transmission of 440- and 560-nm light by the eye’s optics was corrected using an ocular transmission spectrum that was recently measured^33^. This spectrum shows less attenuation of shorter wavelengths than a prior estimate^50^. Use of the recent spectrum yields a 440:560 sensitivity ratio that changes with cumulative light history from 7.5 (95% CI: 5.1-9.0) to 1.1 (95% CI: 0.8-1.4). Use of the prior spectrum yields ratios of 7.9 (95% CI: 5.4-9.6) and 1.4 (95% CI: 1.1-1.9).

#### Electrophysiology

Following dark adaptation (1.5 hours to overnight), mice were anesthetized with Avertin, enucleated, and euthanized via cervical dislocation. The retina was mechanically freed from the retinal pigment epithelium and vitreous humor in Ames medium equilibrated with 95% O_2_/5% CO_2_ (United States Biological or Sigma-Aldrich, supplemented with NaHCO_3_) or a reduced form of Ames medium^22^ (see below). The retina was mounted flat, with rods/cones adhering to a coverslip coated with poly-L-lysine. RGCs faced upward and the tissue was superfused continuously with Ames medium (∼5-8 ml/min). Cells were visualized on an upright microscope using infrared transillumination (850- or 940-nm center wavelength and 30-nm width at half-maximum) and differential interference contrast optics. To identify ipRGCs, a field of view was given <2 s of imaging light (for tdTomato, a 25-nm band centered on 545 nm, ≤ 6 × 10^9^ photons µm^-2^ s^-1^). IpRGCs were exposed by mechanical removal of overlying membranes and dark-adapted for ≥20 minutes prior to measuring light responses. Action spectra studied in this condition reflect melanopsin’s R state, indicating an adequate return of pigment fractions to baseline^7^ (also see above). Temperature was controlled with an inline heater and monitored continuously by a thermistor in the bath solution.

M1 ipRGCs were targeted by their bright fluorescence, small soma diameters, and physiological properties^39,41^. Neurons were only included for analysis if the seal resistance was ≥10 GΩ and the series resistance changed by ≤ 20% over the recording (which lasted as long as 1.5 hrs). Recording pipettes (3-7 MΩ resistance) were generally wrapped with parafilm to reduce capacitance. Series resistance (typically <60 MΩ) was monitored but not compensated. The small size and slow time course of most responses measured (picoamperes flowing over seconds or minutes), combined with the high input resistances of M1 ipRGCs (often >1 GΩ), promote voltage clamp through this high series resistance^41^. Multiclamp 700B amplifiers were used for all experiments. Details concerning signal acquisition and processing are in **Table S1**; filtering was with a 4-pole Bessel filter and sampling always met or exceeded the Nyquist minimum.

#### Electrophysiology Solutions

Ames medium (see above) or a reduced form of Ames medium^22^ was used. The latter contained (in mM) 120 NaCl, 22.6 NaHCO_3_, 3.1 KCl, 0.5 KH_2_PO_4_, 6 glucose, 1.2 CaCl_2_, and 1.2 MgSO_4_. For Ca^2+^ removal (in voltage clamp), the solution was 120 NaCl, 22.6 NaHCO_3_, 3.1 KCl, 0.5 KH_2_PO_4_, 6 glucose, 2.6 MgSO_4_, 1 EGTA, and 0.0005 TTX (the control solution for these experiments also contained 0.0005 TTX). External solutions were equilibrated with carbogen (95% O_2_/5% CO_2_). The antagonists added to block synaptic transmission were (in mM) 3 kynurenate, 0.1 picrotoxin, 0.01 strychnine, and 0.1 D,L-AP4. The perforated-patch mode of electrophysiological recording was used to preserve melanopsin phototransduction, which is sensitive to dialysis^22,62,63^. The internal solution was composed of (in mM) 110 K-Methanesulfonate, 13 NaCl, 2 MgCl_2_ or MgSO_4_, 10 EGTA, 1 CaCl_2_, 10 HEPES, and 0.125-0.25 amphotericin B (pH 7.2 with KOH for a final [K^+^] of 139-140 mM). 0.5% Neurobiotin (Vector Laboratories) was sometimes included in the internal solution.

#### *Ex Vivo* Optical Stimulation

Light from a 75-W Xe arc lamp was filtered to deplete ultraviolet and infrared wavelengths, then sent through additional filters to control wavelength and intensity as needed. Xe itself was used for broadband (white) light because it resembles daylight. To mimic spectral filtering by the eye’s optics, two 81A filters (Cokin Filters) were placed in series. The transmission spectrum of this filter closely resembles that of the mouse ocular transmission spectrum measured by Henriksson and colleagues^50^. A recently measured spectrum^33^ indicates higher transmission of short wavelengths. Use of this recent spectrum would be expected to produce observations that lie between those obtained with Xe and 81A-filtered Xe stimuli.

Light was focused through a 40× objective to produce a spatially uniform disc (350-µm diameter) centered on the soma as epi-illumination. An electromechanical shutter controlled stimulus timing. Stimuli were measured at the site of the preparation using a calibrated radiometer and spectrometer. The spectra of broadband optical stimuli have been published^7^. Light delivered through a 10-nm bandpass filter was assumed to be of the center wavelength. For broader filters, photon flux was calculated from measured spectra.

#### Comparing Monochromatic and Broadband Stimuli

Spectral power distributions were measured for broadband illumination (81A-filtered Xe, referred to as ocular-filtered Xe or Xe_eye_; see above), a 10-nm band centered on 440 nm, and a 10-nm band centered on 560 nm. Measurements were made using a fiber-coupled spectrometer. While the shape of the spectrum is accurate, its amplitude is sensitive to fiber angle. Therefore, the amplitude was scaled according to the following procedure: (1) the 440-nm band’s power was measured with a radiometer; (2) the 440-nm band in the spectrum was scaled such that its integral matched the radiometer’s power reading; and (3) this scale factor was applied to the Xe_eye_ spectrum. Power spectral density was converted to photon flux density (photons µm^-2^ s^-1^ nm^-1^), and the total intensity of Xe_eye_ obtained by integration.

To obtain scale factors that equate Xe_eye_ and monochromatic lights (440 and 560 nm), these illuminants were given to M1 ipRGCs. Perforated-patch, voltage-clamp recordings (-80 mV) were conducted in synaptic antagonists, with heat cycling (see above) to improve dark adaptation. Light was given in flashes (durations of 25-50 ms). Each cell was tested with all three illuminants. Six flashes were given of each illuminant; illuminants were interleaved; and flashes were separated by 90 s for full response recovery in between. Flash intensities were adjusted to obtain peak photocurrents in the linear range (<15 pA)^22,41^. The amplitude of the photocurrent was divided by the flash intensity, using the photon flux densities obtained above and multiplying by the flash duration. This yielded dim-flash sensitivity (S_F_) ratios: Xe_eye_:440 was 0.18 ± 0.02 and Xe_eye_:560 was 1.85 ± 0.26 (6 cells). The 440:560 ratio is given above (10.6 ± 0.6; 6 cells).

To measure pulse I-R relations with Xe_eye_, its intensity was matched to that of 440-nm light (at the retina) using their relative S_F_ values. To measure these relations with Xe, the intensity of this light was scaled to deliver the same total photon count as Xe_eye_. At the completion of the two successive I-R relations, a 140-s, 560-nm pulse (6.4 × 10⁹ photons µm^-2^ s^-1^) was delivered to suppress persistent activity, and the preceding pulse was repeated to measure recovery.

#### Analysis of Intensity-Response Relations

The response examined was pupil constriction, cellular photovoltage, cellular photocurrent, or model activity. Each stimulus was a light pulse. Pulses were given in a series comprising one wavelength, either 440 or 560 nm.

A sigmoid function was fit to I-R relations for pupil constriction, ipRGC photovoltage, ipRGC photocurrent, and model outputs

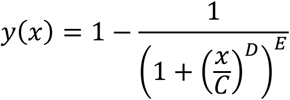

where *C* controls the x-axis position, *D* steepness, and *E* asymmetry. Any response >5% smaller than the maximum response was omitted from the fit.

To quantify the degree to which an I-R relation displayed negative slope, the peak response at the maximum intensity was divided by the largest peak response observed at any intensity (intensities in units of photons μm^-2^). A value <1 indicates a region of negative slope. These values were compared across conditions as needed.

The peak of a response was divided by the photon density of the entire pulse that evoked it, yielding pulse sensitivity. A given pulse sensitivity was plotted against cumulative photon density (summing photons in the considered pulse and all preceding pulses). Comparing between wavelengths (with linear interpolation) yielded a 440:560 pulse sensitivity ratio. The expected ratio following prolonged dark adaptation, when all detectable melanopsin activation is from the R state, is 10.7 (see above). With light history the ratio also reflects activation of melanopsin from the E state (>10.7, because E absorbs 440 nm more effectively) as well as any cumulative effects of light history. Note that ratios compare separate stimulus series. Pulse sensitivity for photocurrent was normalized within each cell before averaging.

Bootstrapping was used to compare 440:560 pulse sensitivity ratios to reference values (e.g. 10.7, noted above, and 1, indicating equal pulse sensitivity for the tested wavelengths). To obtain bootstrap samples, biological replicates (e.g., ipRGCs or mice) were resampled using the Monte Carlo algorithm. Pulse sensitivity and pulse sensitivity ratios were computed for each of 1000 bootstrap resamples.

To estimate spectral convergence expected from response saturation alone (**Figure 2D**), pulse sensitivity ratios were calculated from the sigmoid fit to the short-wavelength I-R relation and a sigmoid of the same shape, shifted by 10.7-fold along the irradiance axis. This shifting simulated a long-wavelength I-R relation of identical shape.

#### Modeling

##### Basic (No Gain Shifting) Model

Melanopsin state photoconversions were modeled numerically as described previously^7^. The states are melanopsin (R), metamelanopsin (M), and extramelanopsin (E):

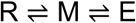

For each state, photon absorption is governed by the extinction coefficient ε (in units of mol^-1^ cm^2^) and a spectral dependency function *A*(λ). Following photon absorption, the probability of isomerization is given by the quantum efficiency (ϕ). Let *F*_*i*_ = ln(10)ϕ_*i*_ε_*i*_*dt* ƒ [*I*(λ)*A*_i_(λ)*d*λ] be the fraction of state *i* that isomerizes due to the photostimulus. *I*(λ) is the light intensity in mol photons cm^-2^ s^-1^ nm^-1^ (higher intensities give faster approaches to equilibrium). The ln(10) term originates with the Beer-Lambert law governing the absorption of light. Light stimuli were delayed by 400 ms to approximate response latency.

The difference equations governing the model are

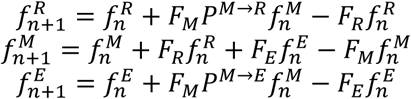

where *f*^*R*^, *f*^*M*^, and *f*^*E*^ are the fractional occupancies of each state and sum to 1. *P*^*M*→*R*^ and *P*^*M*→*E*^ are the probabilities that, if M isomerizes, it transitions to R or E (*P*^*M*→*R*^ = 1 — *P*^*M*→*E*^).

For *A*(λ), Shichida’s broadened photopigment template was used for R (adjusted to fit the ipRGC action spectrum) and M, and Govardovskii’s template for E (see above)^6^. Other model parameters also have been defined biochemically and given by Shichida and colleagues^6^ except for ϕ_*E*_ and *P*^*M*→*R*^ (thus also *P*^*M*→*E*^). Prior work used a ϕ_*E*_ of 0.4, which is intermediate between the values for R and M^6^, and a *P*^*M*→*R*^ (and thus *P*^*M*→*E*^) of 0.5. The model uses a ϕ_*E*_ of 0.65, which improved the prediction of persistent activity. This no-GS base model does not account for light-independent transitions between states^7^.

##### Gain Shifting Model

Adaptation was implemented by a second tier of states, R’, M’, and E’. Transitions of R↔M, M↔E, R’↔M’, M’↔E’ occur through photoisomerization as described above. Conversions between the high-gain (e.g., M) and low-gain (e.g., M’) states are governed by rate constants that vary according to the identity of the state. Melanopsin activities in the no-GS and GS models are *f*^*M*^ and (*f*^*M*^ + 0.038*f*^*M*′^), respectively. The gain of the M’ state was set to 0.038 based on the ratio between the saturated transient response and subsequent persistent activity measured previously^41^. Varying this parameter altered the magnitude of persistent activity (not shown).

The rate constant for entry into the low-gain state (*k*_*MM*_′) was estimated from the decay time constant of the dim-flash response (6.6 s at 35 °C, corresponding to a rate constant of 0.15 s^-1^)^22^. The rate constant of transitioning from the low-gain ground state to its normal-gain form (*k*_*R*_′_*R*_) was 1/22 s^-1^, based on arrestin dissociation from the β2 adrenergic receptor after agonist removal^64^. The same value was used for *k*_*E*_′_*E*_. Varying these rate constants alters the kinetics of flash and step responses, as well as the steady-state magnitude of the step response (not shown). All other rate constants were assumed to be much slower (1/3600 s^-1^). In the table below, *k*_*ij*_ denotes the light-independent rate constant of state *i* to state *j*.

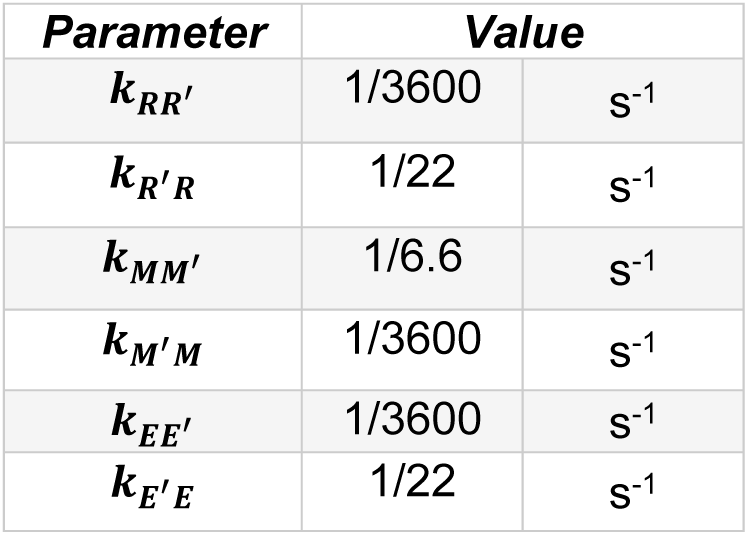

| **Parameter** | **Value** |  |
| --- | --- | --- |
| $k_{RR'}$ | 1/3600 | $\text{s}^{-1}$ |
| $k_{R'R}$ | 1/22 | $\text{s}^{-1}$ |
| $k_{MM'}$ | 1/6.6 | $\text{s}^{-1}$ |
| $k_{M'M}$ | 1/3600 | $\text{s}^{-1}$ |
| $k_{EE'}$ | 1/3600 | $\text{s}^{-1}$ |
| $k_{E'E}$ | 1/22 | $\text{s}^{-1}$ |

The light-dependent difference equations are

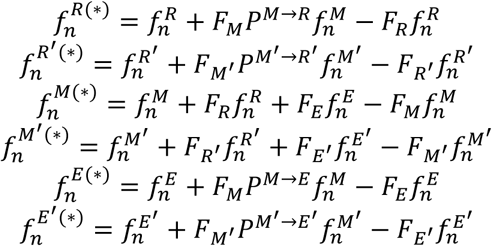

The light-independent difference equations are then applied sequentially within the same timestep.

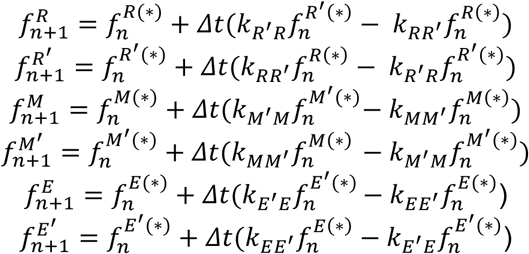

where ^(^*^)^ denotes the intermediate state following a light-dependent transition and prior to a light-independent transition within the timestep *Δt*. At each timestep, total melanopsin is re-normalized to one.

##### Augmented Gain Shifting Model – Loss and Recovery of Melanopsin

Loss and recovery of melanopsin was implemented by a non-photoconvertible state with very low gain, O. The R, M, and E states transition to the O state. The O state recovers to R, the only holopigment state found in dark-adapted ipRGCs^7,60^. The model is initialized at equilibrium, with half of melanopsin in the O state (*k*_*OR*_ = *k*_*RO*_). Rate constants for O recovery to E and M are set to zero in this study but can be adjusted if desired. For transitioning from the E to O state, the rate constant (*k*_*EO*_) was 5/10000 s^-1^. This corresponds roughly to the time elapsed between identification of an ipRGC by tdTomato fluorescence (which produces a large fraction of the E state) and measurement of an action spectrum reflecting only the R state^7^. The rate constants of transitioning from the R, M, and E states to the O state and recovery from O to R were obtained by fitting the model to the empirical I-R relations and photocurrent recordings (see below). The gain of the O state was set to that estimated for bleached tiger salamander rhodopsin, which is ∼1 × 10^-6.5^ (Ref. 65).

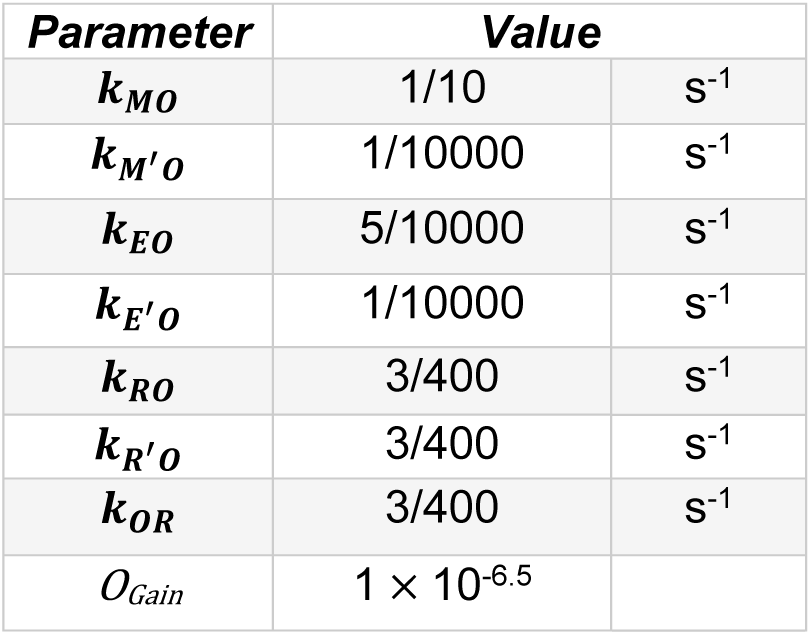

| **Parameter** | **Value** |  |
| --- | --- | --- |
| $k_{MO}$ | 1/10 | $\text{s}^{-1}$ |
| $k_{M'O}$ | 1/10000 | $\text{s}^{-1}$ |
| $k_{EO}$ | 5/10000 | $\text{s}^{-1}$ |
| $k_{E'O}$ | 1/10000 | $\text{s}^{-1}$ |
| $k_{RO}$ | 3/400 | $\text{s}^{-1}$ |
| $k_{R'O}$ | 3/400 | $\text{s}^{-1}$ |
| $k_{OR}$ | 3/400 | $\text{s}^{-1}$ |
| $O_{Gain}$ | $1 \times 10^{-6.5}$ | |

Adding melanopsin loss and recovery to the model recapitulates the graded levels of persistent activity observed in the empirical 440-nm I-R relations. While the GS model predicts that persistent activity reaches a saturated level partway through the I-R relation, a model that includes loss and recovery of melanopsin agrees with the data in predicting graded increases throughout (**Figures 5, 6** and **Figures S4, S5,** and **S9**).

The light-independent difference equations are adjusted to implement exchange with O:

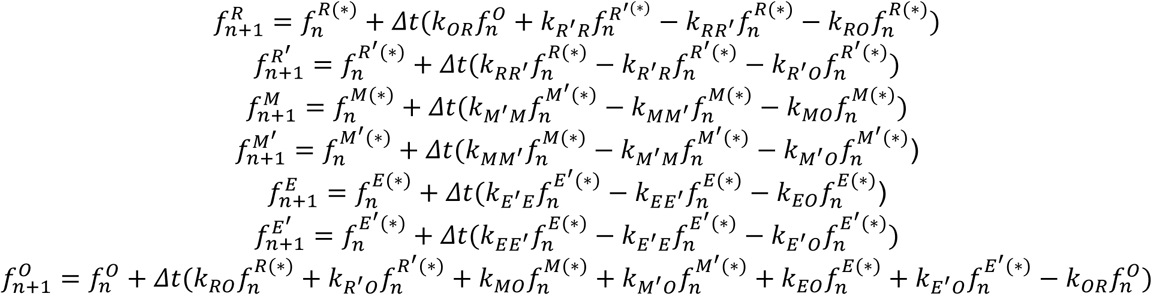

Rate constants for the other light-independent transitions were adjusted to accommodate O.

When delivering pulses of increasing intensity from darkness and measuring photocurrent, the persistent current increased with cumulative light history. Closer inspection reveals that the intensity-current relation appears biphasic, having a shallower slope over the dimmer pulses and a steeper slope over more intense pulses (not shown). This is observed for both 440- and 560-nm light. To approximate it in the aGS model, *k*_*MM*_′ doubles when the M’ fraction exceeds 5%. This feature produces a biphasic increase in the M’ fraction and improves prediction of transient response amplitudes. Note that it does not always make the persistent photocurrent itself biphasic, due to interactions with other mechanisms of adaptation.

| **Parameter** | **Value** |  |
| --- | --- | --- |
| $k_{RR'}$ | 1/3600 | s<sup>-1</sup> |
| $k_{R'R}$ | 1/22 | s<sup>-1</sup> |
| $k_{MM'} (f^{M'} \leq 0.05)$ | 1/83 | s<sup>-1</sup> |
| $k_{MM'} (f^{M'} > 0.05)$ | 2/83 | s<sup>-1</sup> |
| $k_{M'M}$ | 1/833 | s<sup>-1</sup> |
| $k_{EE'}$ | 1/3600 | s<sup>-1</sup> |
| $k_{E'E}$ | 1/22 | s<sup>-1</sup> |

##### Augmented Gain Shifting Model – Negative Regulation

The model was augmented with two negative regulation mechanisms that dynamically adjust the gain of melanopsin’s signaling states. Each regulation mechanism includes a gain control variable that accumulates via leaky integration.

The first gain control (*G*) accumulates from total melanopsin activity (Ψ):

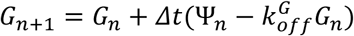

The second gain control (*G*^*M*^) tracks persistent activity and therefore accumulates from melanopsin in the low-gain signaling state, M’:

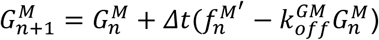

Both gain control variables (*G* and *G*^*M*^) pass through nonlinearities (σ and σ^*M*^) before scaling the gain of the melanopsin signaling states. *G* scales the gain of M, M’, and O, while *G*^*M*^scales the gain of M:

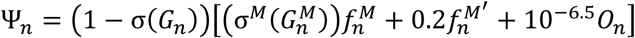

The fixed gain of the M’ state was increased to 0.2 to maintain a realistic ratio between persistent and transient activity in tandem with negative regulation.

Nonlinearities were defined by the equations

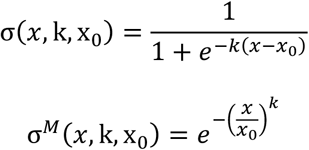

where k controls steepness and x_0_ is the input to reach 50% or 37% gain for σ and σ^*M*^, respectively.

The parameters in the following table were set by fitting the model to empirical I-R relations. Fits were improved with the sigmoid midpoint positioned near zero integrated activity, such that activity near baseline contributes to negative regulation.

| *Parameter* | *Value* |  |
| --- | --- | --- |
| $k_{off}^G$ | 0.037 | s<sup>-1</sup> |
| $k_{off}^{GM}$ | 0.355 | s<sup>-1</sup> |
| $\sigma: (\mathbf{k}, \mathbf{x}_0)$ | 40.0, 0.0008 | |
| $\sigma^M: (\mathbf{k}, \mathbf{x}_0)$ | 1.215, 0.381 | |

Adding negative regulation to the model produces agreement between simulated and empirical I-R relations. Most notably, negative-slope I-R relations emerge in response to simulated short-wavelength pulses (**Figure 6** and **Figure S4**). This is because short-wavelength light drives significant graded persistent activity. This reduces, through *G*^*M*^, the gain of the high-gain signaling state (M). This outcome also depends on loss and recovery of melanopsin. In models lacking this feature but equipped with negative regulation, persistent activity and negative regulation saturate earlier in the I-R relation, resulting in the lack of a negative slope region (**Figure S6**).

Negative regulation also allows the model to capture features of the response kinetics. During simulated high-intensity light stimulation, high melanopsin activity recruits negative regulation through *G*, which reduces gain as transient responses rise. Consistent with empirical data, simulated transient responses to brighter light peak sooner and decay more quickly (**Figure 6** and **Figure S5**). After light offset, negative regulation decays slowly enough to reduce the gain of ongoing persistent activity, producing inflections resembling those observed empirically (**Figures 6, S5,** and **S6**).

Stimuli for simulated I-R relations (**Figures 6**, **S4**, and **S9**) mimic the stimulus set presented during voltage clamp experiments depicted in **Figures 2, 3** and **Figures S2** and **S8**.

##### Augmented Gain Shifting Model – Parameter Fitting

Parameters not estimated directly from measurements were fit to I-R relations (all cells) and photocurrents (a representative cell for each stimulus) for stimuli in **Figures S4** and **S9**. Loss functions computed the sum of squared errors between normalized photocurrents and model predictions, and between cellular and model I-R relations (transient and persistent response amplitudes, normalized to the peak transient response). Fitting consisted of loss-minimization through hand-tuning and algorithmic optimization (differential evolution, basin-hopping, and L-BFGS-B)^66-68^. The aGS model accurately predicted responses to several stimuli that were not involved in parameter fitting (**Figures S5** and **S7**).

To determine whether tuning of the parameters shared between aGS and GS (***k***_***MM***_′, ***k***_***M***_′_***M***_, and M’ gain) could alone account for the aGS model’s improved performance, two GS model variants were evaluated. In the first, GS took on the aGS parameter values. In the second, these parameters were optimized using the L-BFGS-B algorithm to improve predicted I-R relations. Compared to aGS, all GS variants fit the data to a lesser extent: the mean loss for aGS was 0.64, for GS 1.94, for GS adopting aGS parameters 9.89, and for GS with optimized parameters 1.60 (**Figure S10**).

##### Note on Model Design and Performance

Model parameters are not meant to recapitulate specific adaptational mechanisms within melanopsin phototransduction, of which there are many^17,18,20-23,44,46-49^. The goal was to find a minimum number of parameters that can explain the experimental observations. Parameters are biologically inspired but each is likely to reflect a combination of actual mechanisms. The combination of parameter values is also unlikely to be a unique solution.

Models reproduced data from the present work as well as from a published study (**Figure S7**)^7^. This includes accounting for melanopsin fractions in darkness (all melanopsin holopigment is in the R state following a prolonged period in darkness^60^) and after illumination (large and enduring fractions of the M and E states for short- and long-wavelength illumination, respectively^7^).

##### Accessing the Models

Models are available on GitHub (https://github.com/MichaelDo-Lab/melanopsin-model). Users can run models through a graphical user interface, a Jupyter notebook, or from source code.

#### Quantification, statistics, and data availability

Data were analyzed in Clampfit (part of pClamp), DeepLabCut, Igor Pro, MATLAB, and Python. Non-parametric statistics were used as described in the main text, unless otherwise noted. See **Table S1** for methodological details and **Table S2** for statistical results. All data reported in this paper will be shared upon request.

---

**Figure S1.**
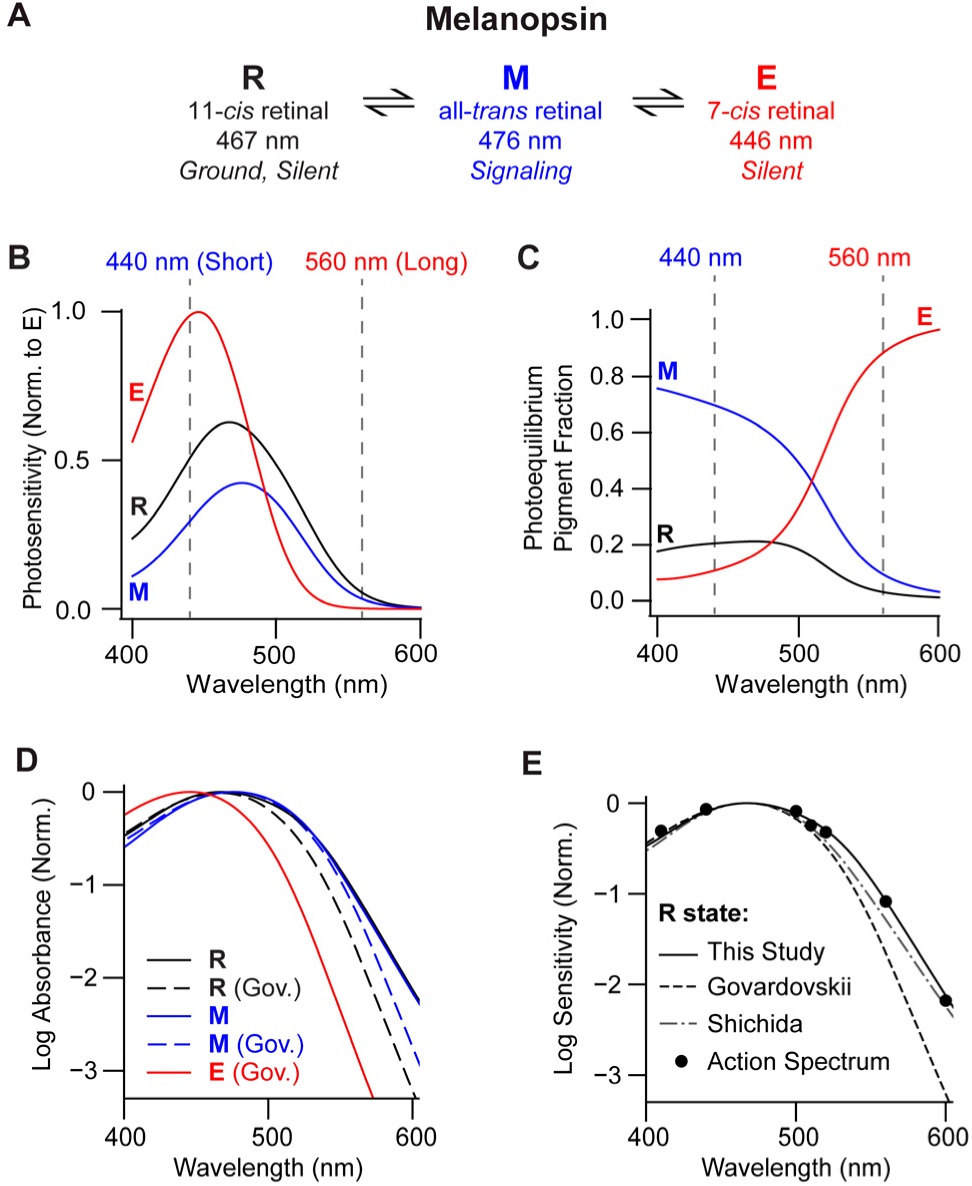
Melanopsin’s absorption and action spectra, related to Figures 1-6. **A.** State diagram for mouse melanopsin holopigment (opsin bound to chromophore) with each conformational state’s chromophore isomer and wavelength of peak sensitivity (λ_max_), from the experiments of Shichida and colleagues on purified mouse melanopsin^6^. R is the only state found in darkness^7,60^, M is the signaling state, and E appears electrically silent^7^. Light is understood to drive interconversions between R and M, and between M and E. Electrophysiological measurements indicate that these states exist in ipRGCs of both mice and macaques^7-9^. Because the M state has high thermodynamic stability, it remains active after illumination ceases, driving minutes-long persistent activity^7,8,42^. **B.** Photosensitivities of melanopsin states, normalized to the maximum of E. The curve for R is a broadened spectral template (e.g. a Govardovskii nomogram^4^ with custom parameters; see **Methods**) fit to the action spectrum of dark-adapted mouse ipRGCs^7^. The curve for M is from the same equation, with parameters fit to purified mouse melanopsin^6^. The curve for E is a standard Govardovskii nomogram with λ_max_ = 446 nm (the value for purified mouse melanopsin^6^). The two principal wavelengths used in this study are marked: 440 nm is absorbed well by all states and 560 nm poorly. Photosensitivities are updated according to best-fit model parameters obtained in the present study (Figure 6 and **Methods**). **C.** Fractional occupancies of mouse melanopsin states at photoequilibrium, as a function of wavelength. The M state, having the longest λ_max_, has high and low occupancy at short and long wavelengths, respectively. The E state, having the shortest λ_max_, follows the opposite pattern. The R state, with an intermediate λ_max_, has low occupancy at every wavelength. Thus, ipRGC activation can be toggled between high and low using acute illumination with short and long wavelengths, respectively. The two principal wavelengths used in this study are marked. 440 nm (short) produces high occupancy of the M state and thus high activation. It also produces R and E state occupancies resembling those under broadband illumination like daylight^7^. 560 nm (long) is effective for deactivation; it produces high occupancy of the E state, and is absorbed well enough to produce photoequilibrium within minutes at light intensities that are readily produced^7^. **D.** *S*pectral templates used in this study (solid lines). The E state uses the Govardovskii template (Gov.). The M state uses the "Shichida" template and fit parameters^6^. The R state uses the Shichida template with parameters fit to the dark-adapted ipRGC action spectrum^7^. Govardovskii fits to the M and R states are also given for comparison (dashed lines)^4^. The log axis emphasizes template shapes in the long-wavelength region. R and M spectral templates overlap closely and are broader than their Govardovskii templates. **E.** Overlay of the action spectrum of dark-adapted ipRGCs (points)^7^ and its Shichida fit (solid line), a Shichida fit to the absorption spectrum of purified melanopsin in the R state (dashed-dot line)^6^, and a Govardovskii template for the R state (dashed line; λ_max_ = 467 nm)^4^.

**Figure S2.**
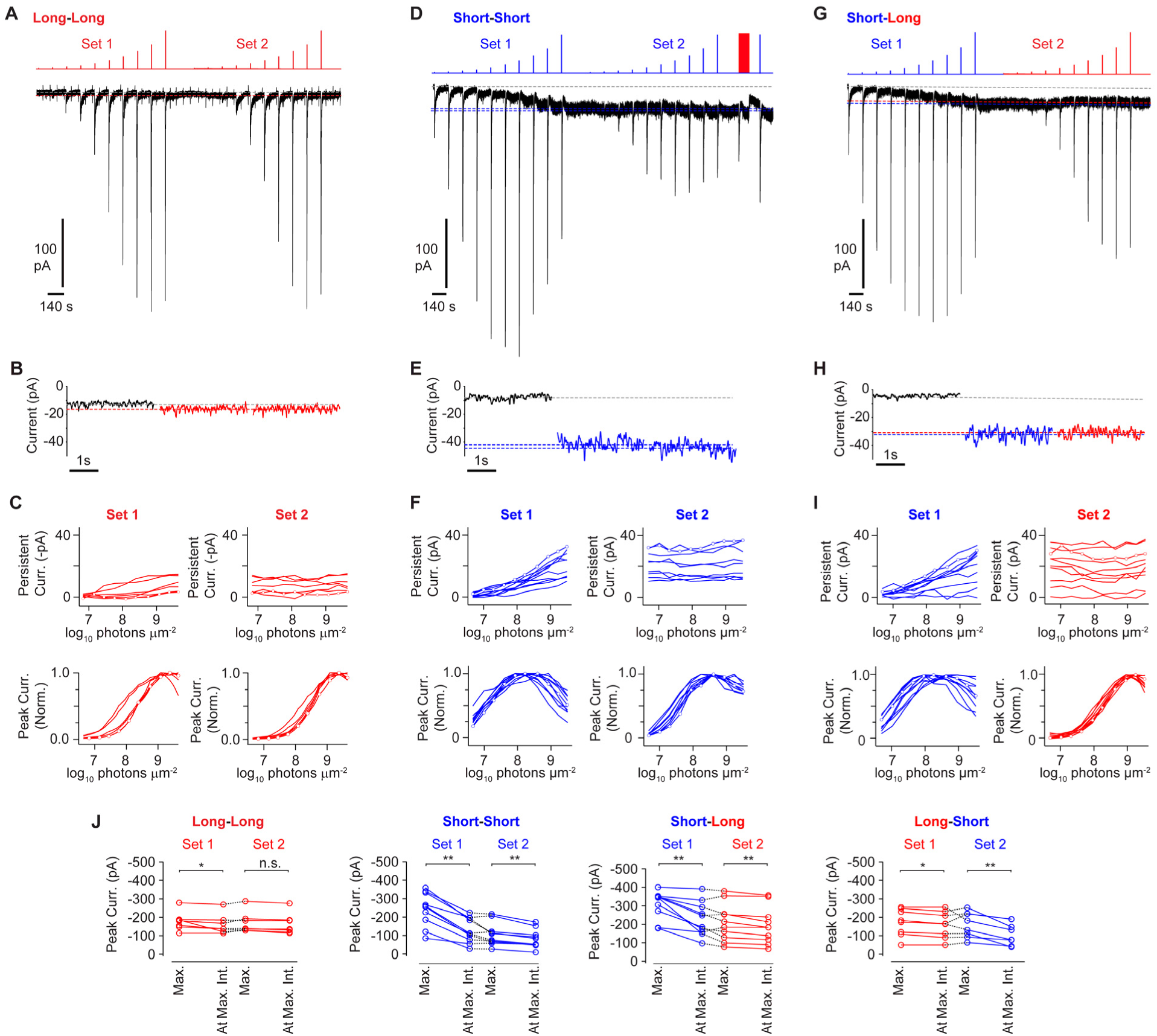
Spectral sensitivity of melanopsin phototransduction current, related to Figures 2 and 3. **A.** Example of a cell given two consecutive sets of intensifying, long-wavelength pulses. Persistent current rises modestly across the first set and remains largely stable across the second. The transient peak amplitude increases within each set and remains near saturation. In all example cells, 10-mV hyperpolarizing steps were given periodically to probe recording parameters and capacitance transients are sometimes observable. **B.** Excerpts of persistent activity in panel **A** at baseline (*left*), after the first set (*middle*), and after the second set (*right*). **C.** I-R relations for persistent current (*top*) and peak transient current (*bottom*). Transient activity is normalized for ease of visualization. Open circles designate the example cell. **D.** Example of a cell (same as in Figure 2A) that was given two consecutive sets of intensifying, short-wavelength pulses. Persistent current rises across the first set and remains largely stable across the second. The transient peak amplitude increases and then decreases within each set, giving two intensity-response (I-R) relations with regions of negative slope. Following the second set, a pulse of long-wavelength light suppressed persistent activity and a subsequent short-wavelength pulse revealed sensitization (observed in 3 of 3 additional cells tested with the same protocol). Dotted lines mark the pre-stimulus baseline (gray) and the persistent response magnitude following the first and second sets (blue). **E.** Same as panel **B**, but for the example cell in **D**. **F.** Same as panel **C**, but for cells exposed to the short-short stimulus set. **G.** Example of a cell given short- and then long-wavelength pulses. Dotted lines mark the pre-stimulus baseline (gray) and the persistent response magnitude following the first (blue) and second (red) sets. **H.** Same as panel **B**, but for the example cell in **G**. **I.** Same as panel **C**, but for cells exposed to the short-long stimulus set. **J.** Maximum peak transient current and peak transient current at maximum intensity are shown for long-long, short-short, short-long, and long-short pulse series. See **Table S1** for additional methodological details and **Table S2** for statistical results (* p < 0.05, ** p < 0.01, *** p < 0.001). Antagonists of synaptic transmission were included in all experiments.

**Figure S3.**
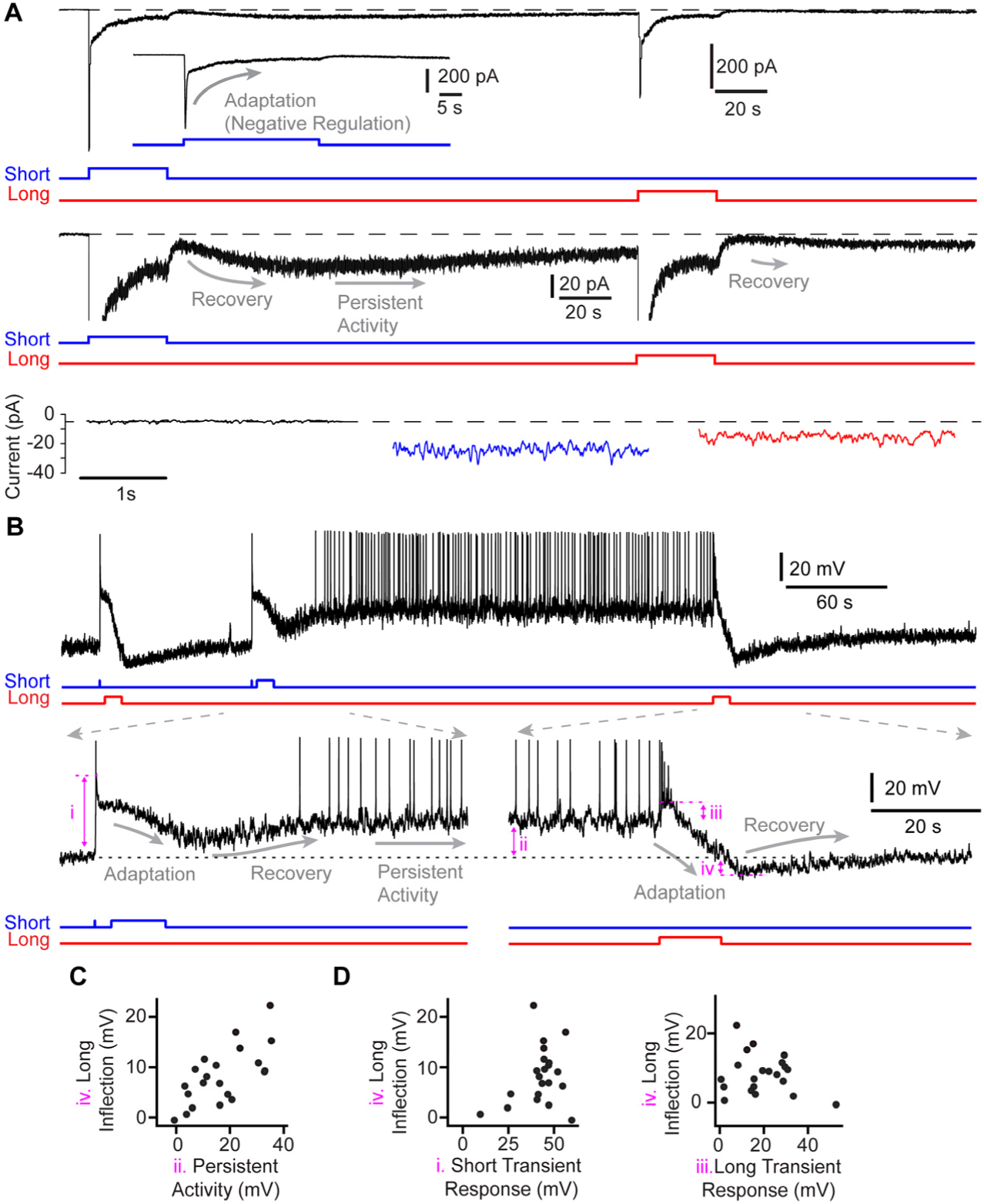
Interaction of persistent activity and adaptation in ipRGCs, related to Figures 3 and 4. **A.** The membrane voltage is clamped (-80 mV) to suppress voltage-gated ion channels and visualize the current driven by melanopsin phototransduction. *Top,* Short-wavelength induction light (blue stimulus monitor) evokes an intrinsic, persistent photocurrent in ipRGCs whose magnitude is attenuated by long-wavelength suppression light (long; red stimulus monitor). *Middle,* The photocurrent on an expanded ordinate. The induction light evokes a large transient current that, even during the pulse, decays toward baseline, which is a sign of adaptation in phototransduction. Adaptation reverses gradually and, as it does, a relatively stable plateau of persistent activity (evident as current and noise) appears. The result is a post-illumination inflection in the current (observed in 7 of 7 additional cells). The suppression light also drives adaptation that decays to reveal attenuated persistent activity, also with a post-illumination inflection (observed in 6 of 7 additional cells). These dynamics indicate that adaptation acts on persistent activity in melanopsin phototransduction. *Bottom,* Excerpts of current in darkness (zeroed), after the short-wavelength pulse (persistent current), and after the long-wavelength pulse (following suppression of persistent current). **B.** Negative regulation of persistent activity is also evident at the level of membrane voltage. Different cell from panel **A**. *Top*, from left to right: A short-wavelength induction flash (blue) would induce persistent activity if not for a subsequent, long-wavelength suppression pulse (red). If that subsequent pulse is short-wavelength instead, persistent activity is evident. Persistent activity is suppressed by a later, long-wavelength pulse. Note that adaptation driven by both short- and long-wavelength pulses is initially strong, cutting into persistent activity and creating an inflection (observed in 16 of 20 cells at 23 °C, including the example cell, and 3 of 3 cells at 35 °C). *Bottom,* these dynamics are more evident on expanded time axes. Measured response parameters are marked in magenta; note that *i* concerns the peak subthreshold voltage (i.e., omitting any action potentials). **C.** Hyperpolarizing inflections after the suppression pulse are correlated with the amount of persistent activity, suggesting that persistent activity drives adaptation (evaluated in the same 20 cells at 23 °C). The larger the persistent activity, the greater the adaptation it drives, and the greater the hyperpolarization that appears once persistent activity is suppressed. In other words, graded persistent activity drives graded adaptation. **D.** No correlations are evident between the magnitude of the transient response to light and the post-transient inflection, which suggests that adaptation is not as strongly driven by the light that induces and/or suppresses persistent activity. Collectively, these analyses indicate that adaptation acts on and is produced by persistent activity. See **Table S1** for additional methodological details and **Table S2** for statistical results. Antagonists of synaptic transmission were included in all experiments.

**Figure S4.**
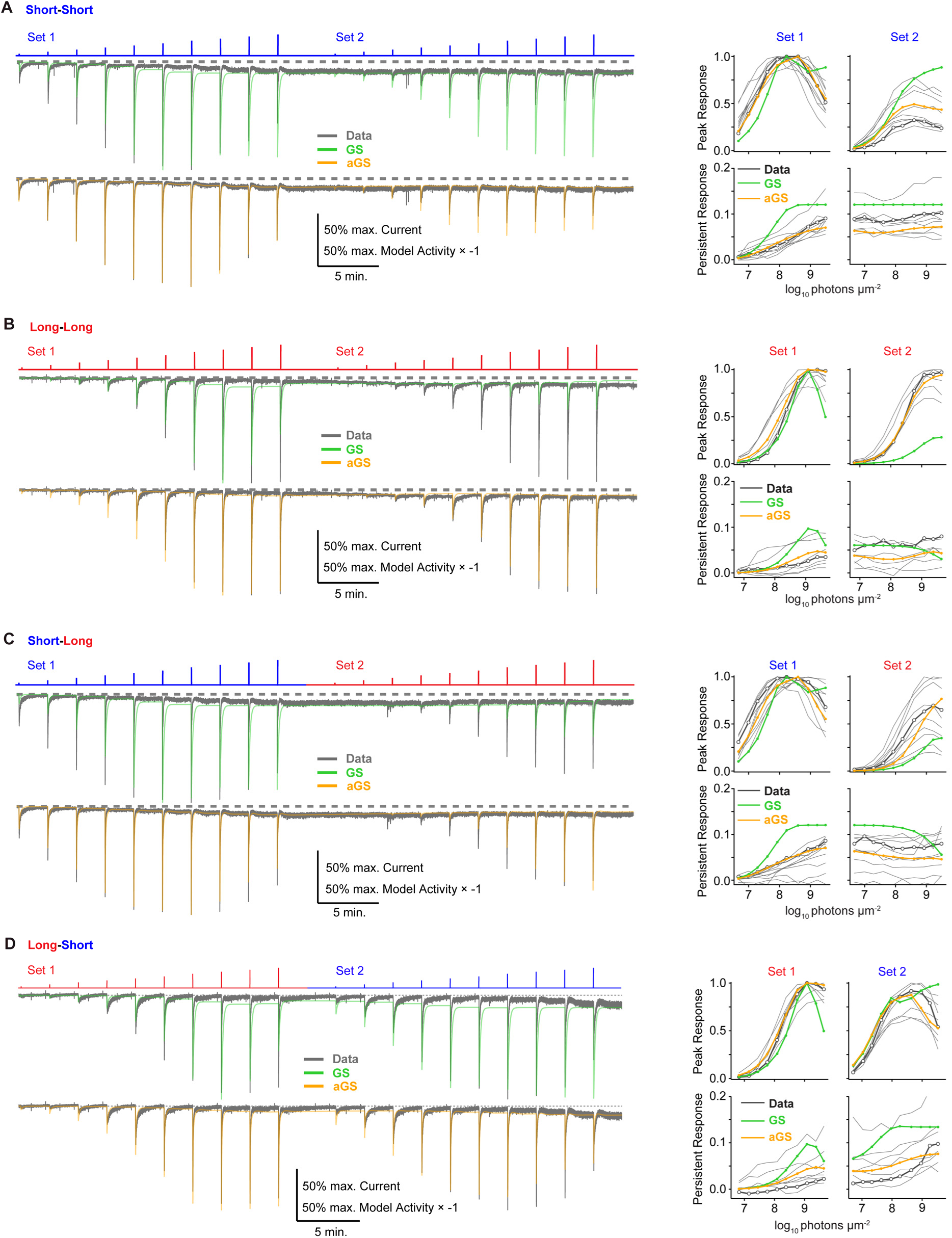
Model predictions of intensity-response relations, related to Figures 5 and 6. **A.** *Left,* Responses of the gain shifting (GS; above, in green) and augmented gain shifting (aGS; below, in orange) models compared to an example ipRGC (gray traces; same trace is compared to each model’s output) given the short-short stimulus (compare to **Figure S2**). Stimulus monitor is at the top. *Right,* I-R relations for GS (green) and aGS (orange) compared with the sample of ipRGCs (gray lines) and example cell from the left panel (black line with open circles). Model and cellular responses are normalized to their peak transient responses, and model responses are inverted for comparison to photocurrents. **B.** Identical to panel **A**, except for the long-long sets. **C.** Identical to panel **A**, except for the short-long sets. **D.** Identical to panel **A**, except for the long-short sets.

**Figure S5.**
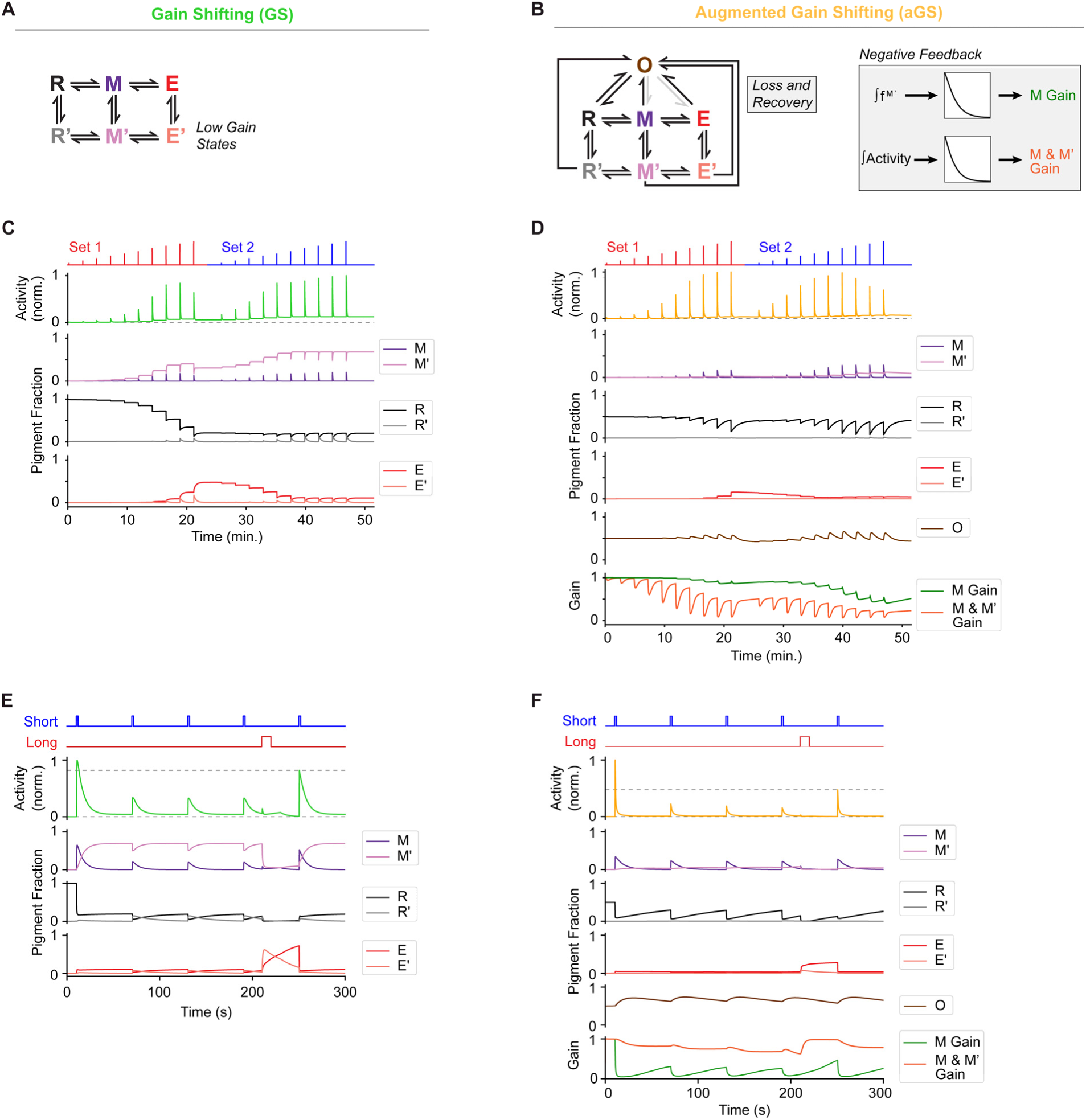
Comparison of gain shifting models with and without augmentation, related to Figures 5 and 6. **A.** Schematic of the Gain Shifting (GS) model. States are color-coded to correspond to traces in panels **C** and **E**. **B.** Schematic of the augmented-GS (aGS) model. *Left*, State transition*s*. *Right*, Negative regulation mechanisms. States and negative feedback mechanisms are color-coded to match traces in panels **D** and **F**. **C.** Melanopsin activity and photopigment fractions predicted by the GS model in response to the long-short stimulus of **Figure S2G** (red and blue, respectively; stimulus monitor at the top). **D.** Melanopsin activity, photopigment fractions, and signaling gains predicted by the aGS model in response to the long-short stimulus of **Figure S2G**. **E.** As in panel **C**, but for the stimulus of Figure 3E consisting of saturating short-wavelength pulses (blue stimulus monitor) and a long-wavelength step to suppress persistent activity (red stimulus monitor). **F.** As in panel **E**, but for the aGS model. Consistent with cellular responses (Figure 3E), aGS predicts less recovery of the saturated response after suppression of persistent activity. Cellular responses to these stimuli were not used for model fitting.

**Figure S6.**
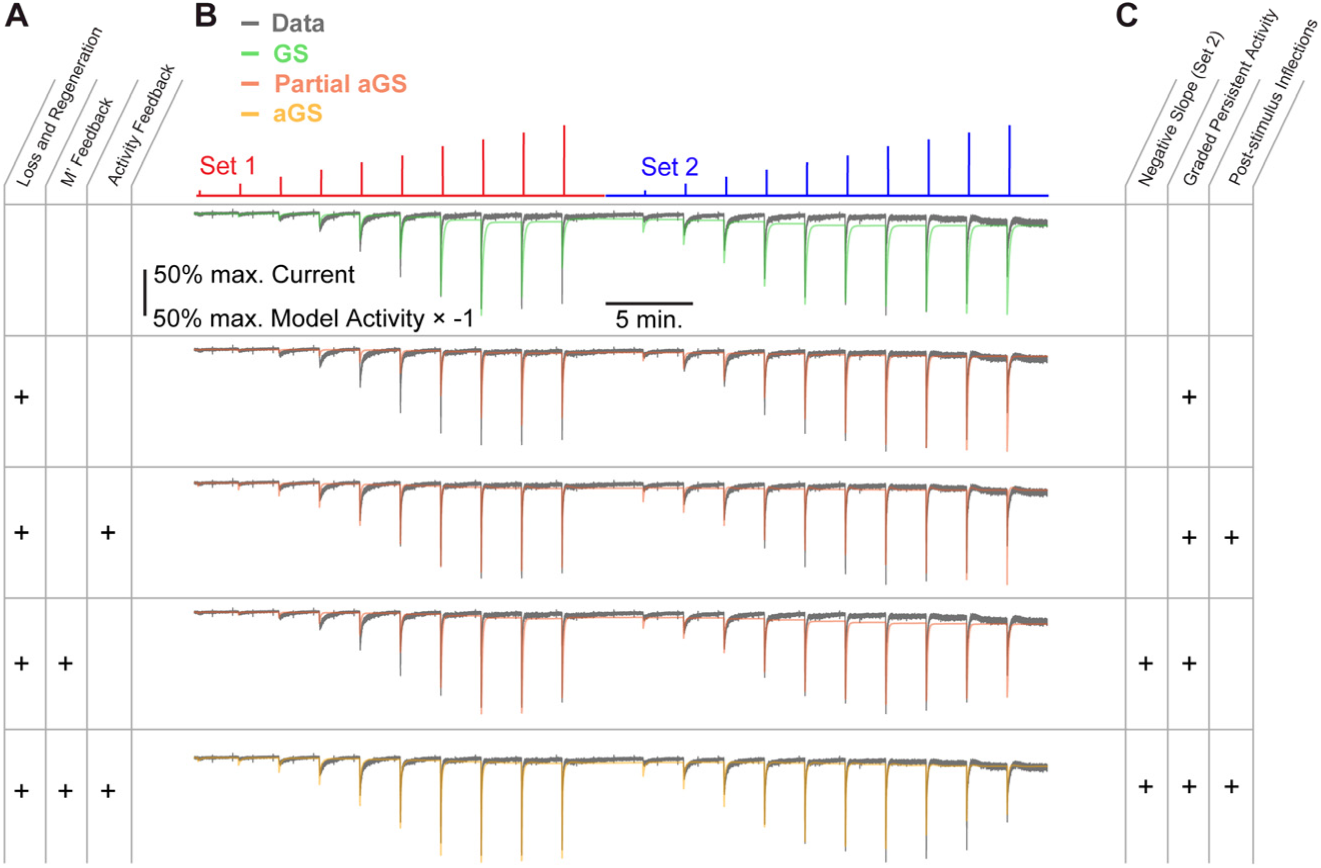
Consequences of individual augmentations of the gain shifting model, related to Figures 5 and 6. **A.** The three available augmentations. “+” designates augmentations added to the model. Rows correspond to the simulations in panel **B**. See **Methods** for additional details. **B.** Melanopsin activity simulated by different models (green: GS model; red: hybrid between GS and aGS; orange: aGS model) compared to an example ipRGC (gray trace, repeated for comparison to each model). The stimulus is a set of intensifying long-wavelength pulses (red stimulus monitor), followed by a set of intensity-matched short-wavelength pulses (blue stimulus monitor; this stimulus set was presented for the experiments of Figure 3A). **C.** Features observed in the responses of real ipRGCs that are captured by each simulation.

**Figure S7.**
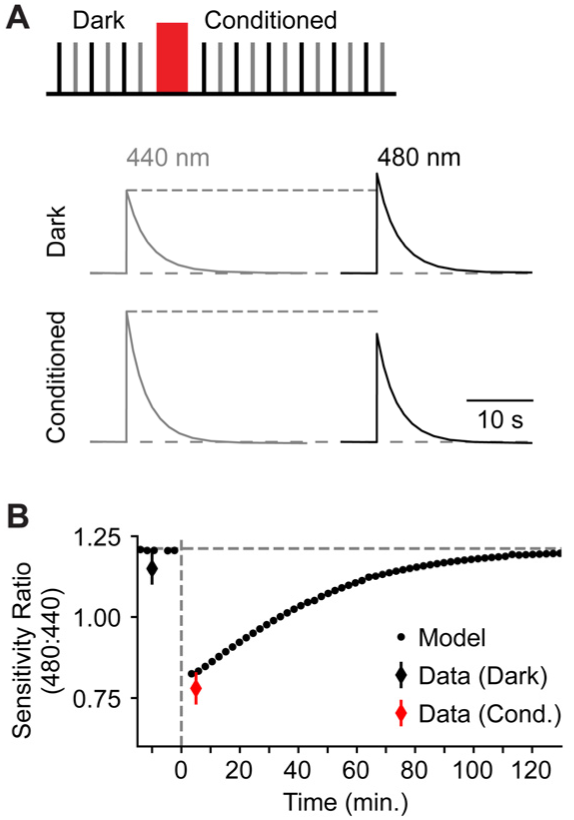
Comparing model predictions to cellular shifts in spectral sensitivity following long-wavelength conditioning light, related to Figures 5 and 6. **A.** *Top:* Protocol for measuring sensitivity before and after a long-wavelength conditioning step (red bar). Black and gray lines in the stimulus monitor indicate 480- and 440-nm flashes, respectively. Sensitivity is higher to 480- or 440-nm light if all melanopsin is in the R or E state, respectively^7^. Long-wavelength conditioning light enriches for the E state (**Figure S1**). *Bottom:* Sample aGS model responses to 440-nm and 480-nm flashes before (‘Dark’) and after long-wavelength conditioning light (‘Conditioned’). **B.** 480:440 sensitivity ratios before and after long-wavelength conditioning light (vertical dashed line at t = 0). Each point is the ratio computed from a pair of flashes. Overlaid data are 480:440 sensitivity ratios measured previously from ipRGCs (n = 9 cells, mean ± SD)^7^. 440- and 480-nm flashes were 50 ms and delivered 10^5.7^ photons µm^-2^. Long-wavelength conditioning: 560-nm, 30 s of 10^9.3^ photons μm^-2^ s^-1^. The model was not fit to these experimental data.

**Figure S8.**
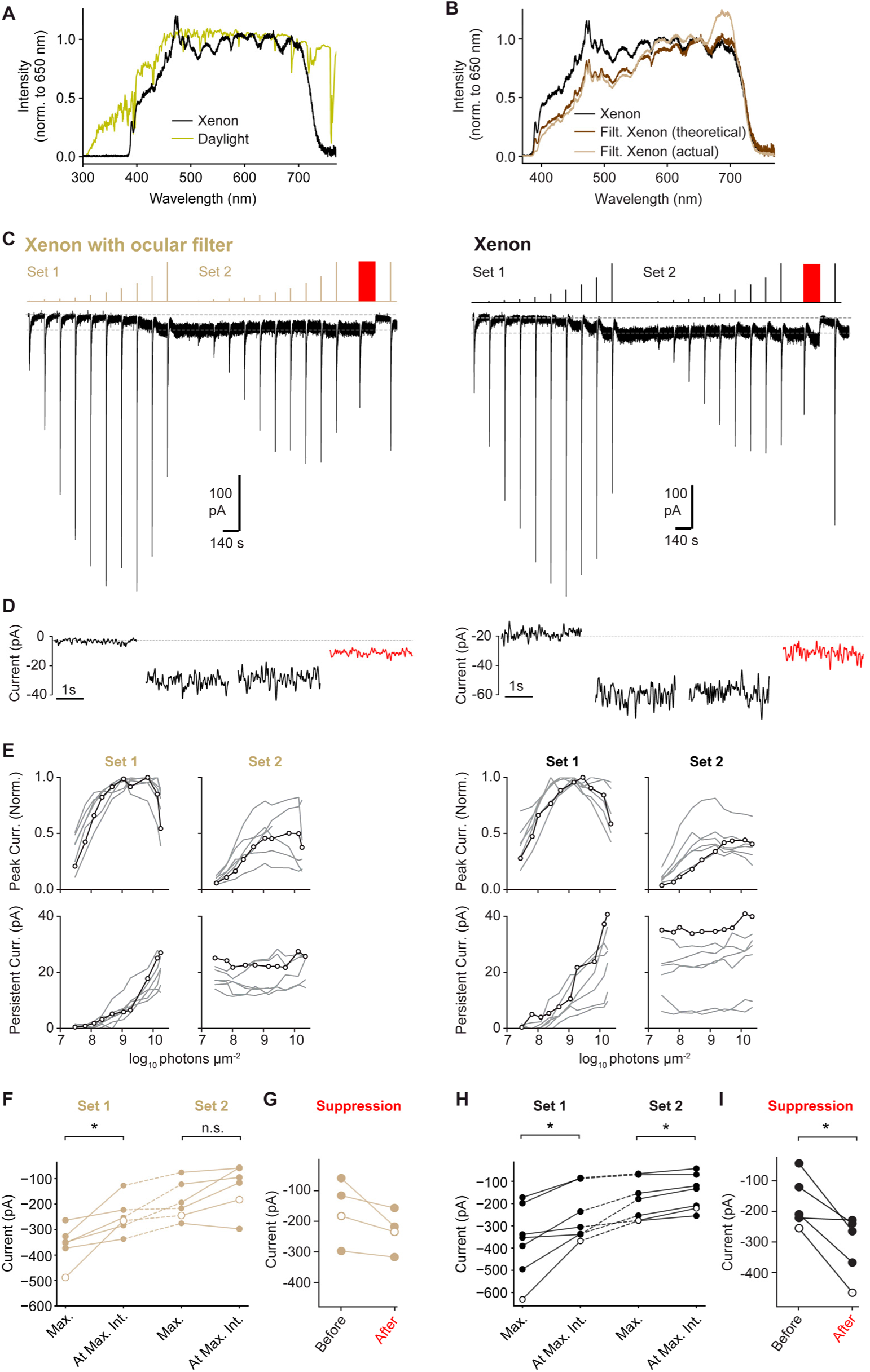
Photocurrent intensity-response relations for broadband (white) light, related to Figures 2 and 3. **A.** Normalized spectrum of daylight (G173 standard^69^; yellow trace) and xenon light (black trace). **B.** Normalized spectra of xenon light (black) and light from the same xenon source after simulated transmission through the ocular optics of the mouse^50^ (brown trace; ‘theoretical’) or measured transmission through an optical filter that mimics these optics (gold trace; ‘actual’; see **Methods**). **C.** *Left:* A voltage-clamped ipRGC (-80 mV) was given two consecutive sets of intensifying, ocular-filtered xenon pulses (stimulus monitor at the top). Following set 2, a pulse of long-wavelength light (red) was given to suppress persistent activity and a subsequent stimulation pulse was given to test for sensitization. *Right:* Same as on the left, but for unfiltered xenon light. Dotted lines mark the pre-stimulus baseline (gray). 10-mV hyperpolarizing steps were given periodically to probe recording parameters and capacitance transients are sometimes observable. **D.** From left to right, excerpts of persistent activity for the two cells in panel **C** at baseline, after set 1, after set 2, and after long-wavelength light given to suppress persistent activity. **E.** Intensity–response (I–R) relations for peak transient (top) and persistent (bottom) melanopsin photocurrents elicited by ocular-filtered and unfiltered xenon light (left and right clusters, respectively). Example cells in **C** are represented by open circles. **F.** Amplitudes of the maximum peak photocurrent evoked and of the peak photocurrent evoked by the maximum pulse intensity for set 1 and set 2 of ocular-filtered xenon. **G.** Peak photocurrent amplitude after long-wavelength illumination, for ocular-filtered xenon. **H.** As in **F** but for unfiltered xenon. **I.** As in **G** but for unfiltered xenon. See **Table S1** for additional methodological details and **Table S2** for statistical results (* p < 0.05, ** p < 0.01, *** p < 0.001). Antagonists of synaptic transmission were included in all experiments.

**Figure S9.**
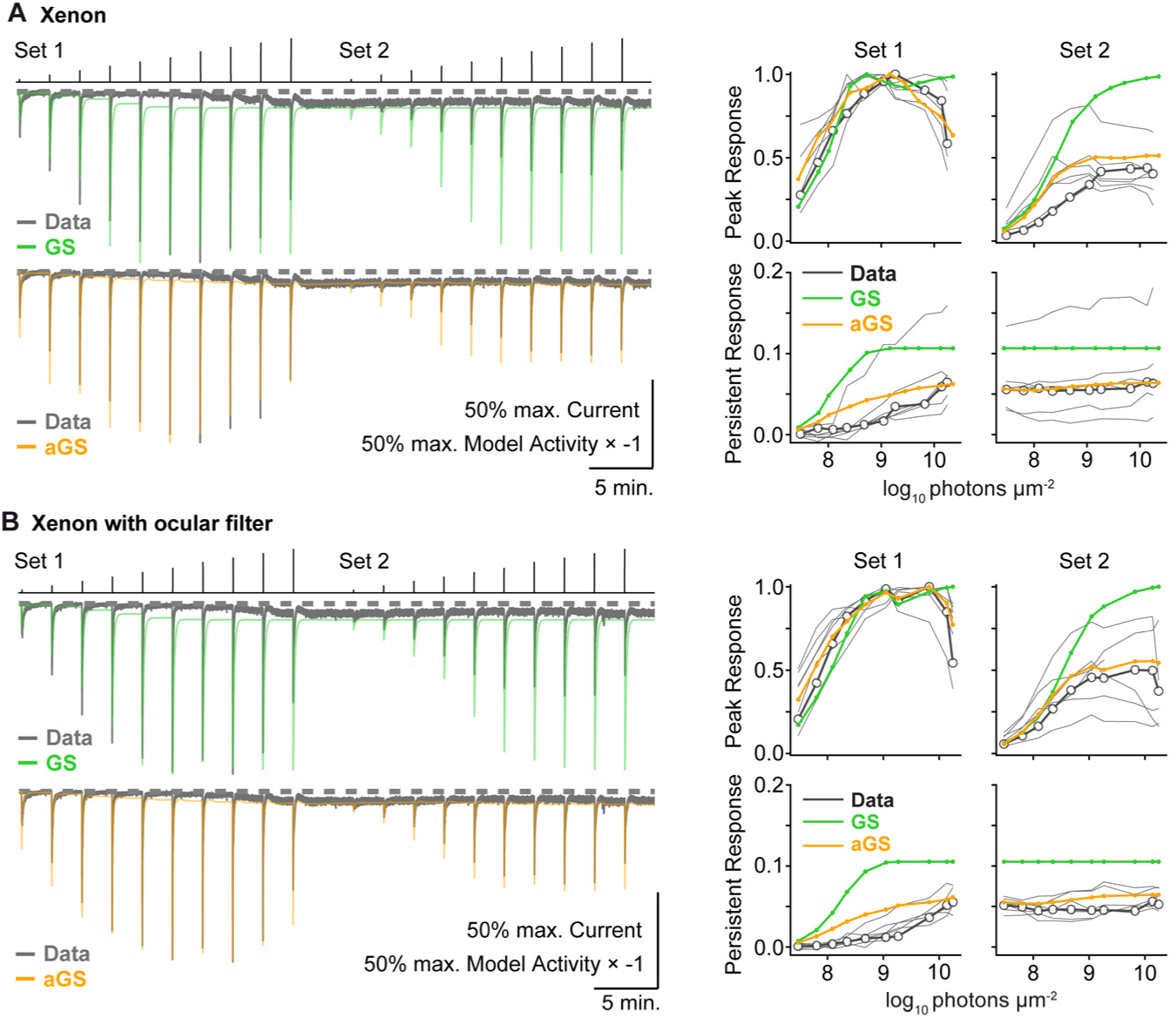
Model responses to broadband illumination, related to Figures 5 and 6. **A.** *Left:* Responses of the gain shifting (GS; green) model, the augmented gain shifting (aGS; orange) model, and an example ipRGC (gray traces) stimulated with xenon pulses. The same gray trace is repeated for comparison to each model. Stimulus monitor is at the top. *Right:* I-R relations, normalized to the maximum transient response of each cell. Open circles and thicker lines mark the example cell. **B.** Same as panel **A** but for ocular-filtered xenon pulses (see **Figure S8B**).

**Figure S10.**
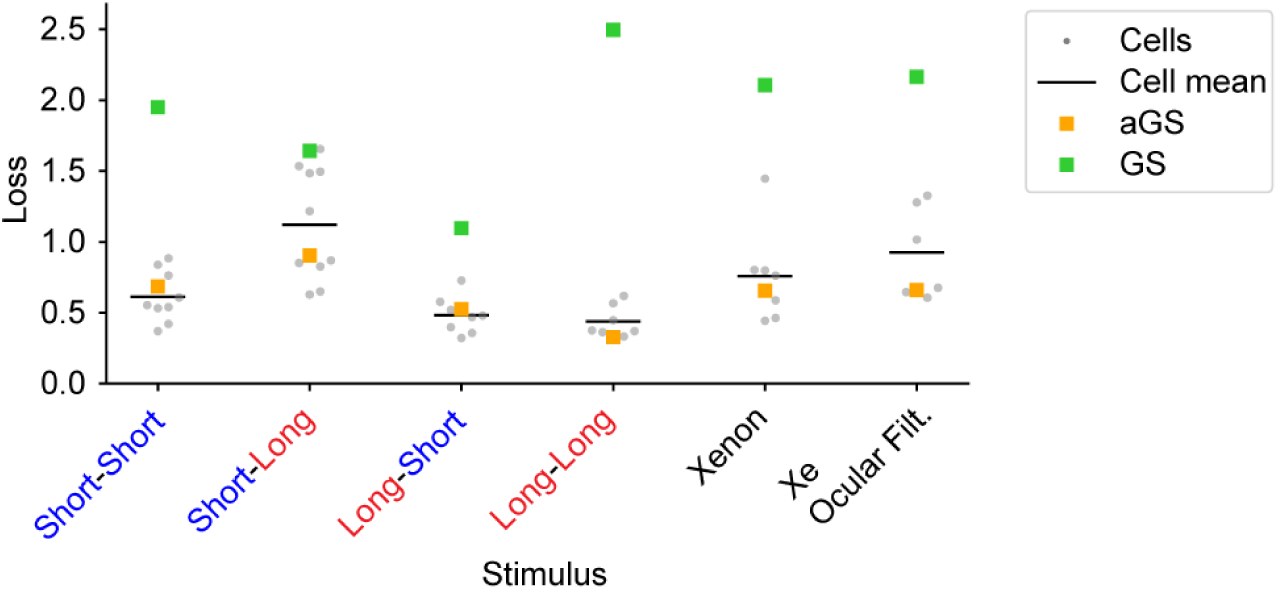
Model prediction benchmarks, related to Figures 5 and 6. Loss (mean squared error between predicted and cellular I-R relations after fitting the aGS model) with predictions provided by the GS model (green squares), aGS model (orange squares), or individual cells (gray dots). Black bars indicate the mean loss of cellular predictions.

**Table S1.** Methods details. The table does not include stimulus intensities for evoking dim-flash responses. These stimuli are calibrated to drive each M1 ipRGC’s melanopsin phototransduction within its linear range, which is typically ≤10 pA^22,41^.

| Figure | Wavelength (nm) | Duration (s) | Retinal Intensity ($\log_{10}$ ph./$\mu\text{m}^2/\text{s}$) | Processing | Measurement |
| --- | --- | --- | --- | --- | --- |
| **1A-E** | 440<br>560 | 1<br>1 | 4.72-8.95<br>4.93-9.14 | Acquisition: 30 Hz sampling. Additional 1 Hz (30 frame) median filter. DeepLabCut was used for point tracking, and points with a likelihood score <0.6 were discarded. | Peak constriction: minimum pupil diameter during the stimulus and inter-stimulus interval, baseline subtracted (1-s window immediately prior to the stimulus). |
| **1F-I** | 440<br>560 | 1<br>1 | 4.72-9.49<br>4.92-9.68 | Acquisition: 10 kHz low-pass filter, 50 kHz sampling. Additional 2 Hz low-pass filter for responses of small and intermediate magnitudes (and thus slow kinetics<sup>22,41</sup>). No additional filter for large responses. | To remove the influence of action potentials, transient subthreshold responses were measured from fits of low-pass filtered traces. Larger responses that lacked action potentials (due to depolarization block<sup>42</sup>) were measured directly (100-ms window around the peak). Baselines were subtracted (1-s window prior to each stimulus lacking action potentials). |
| **2A-D**<br>**3A-D** |  |  |  |  |  |
| Stimulation | 440<br>560 | 1<br>1 | 6.70-9.52<br>6.82-9.64 | Acquisition: 4 kHz low-pass filtering, 10 kHz sampling. Additional low-pass filter: 10 Hz for persistent responses, 2-10 Hz for transient responses. | Transient response: 100-ms window centered on the response peak. Persistent response: 130-140 s after light stimulation. Baseline subtracted (10-s window prior to the first light stimulus).<br>Suppression pulse analysis (Figure 3A):<br>Transient peak: 100-ms window centered on the response peak.<br>Steady response: 10-s window prior to illumination ceasing (130-140 s after light stimulation). |
| Suppression | 560 | 140 | 9.64 |  |  |
| Post-Suppression | 440 | 1 | 9.52 |  |  |
| **3E-H** |  |  |  |  |  |
| Short<br>Long | 440<br>577 | 2<br>2 | 9.23<br>10.60 | Acquisition: 4 kHz low-pass filtering, 10 kHz sampling. Additional low-pass filter $\geq 10$ Hz. | Transient response: average current in a 50-ms window centered on the peak.<br>Persistent response: average current in a 5-s window immediately prior to the saturating pulse. |
| **3E-H**<br>Suppression<br>Induction | 577 or >575<br>440 | 10 or 30<br>10 | 10.60 or 10.25<br>9.23 |  |  |
| **4A-B**<br>Induction<br>Suppression | 440<br>560 | 2<br>40 | 9.56<br>9.69 | Acquisition: 4 kHz low-pass filtering, 10 kHz sampling. Additional 10 Hz low-pass filter for persistent responses. | Persistent response: 10-s window taken 3 min after light stimulation or 140 s after solution exchange. Baseline: 10-s window prior to flash onset. |
| **4C** | 560 | 0.05 | 8.13-9.69 | Acquisition: 4 kHz low-pass filter, 10 kHz sampling. Additional 10 Hz low-pass filter for persistent responses. | Dim-flash response: average current in a 400-ms window centered on the response peak. Dim-flash sensitivity: dim-flash response amplitude (in pA) divided by flash intensity. |
| **5B** | 480 | 0.1 | 7-10 | Model predictions with and without gain shifting. 1-2 kHz time resolution. | N/A |
| **5C** | 480 | 0.2<br>120<br>0.2 | 7.5<br>3-8<br>7.5 | Model predictions with and without gain shifting. 200 Hz time resolution. | N/A |
| **5D**<br>Stimulation<br>Suppression | 440<br>560 | 2<br>20 | 9<br>10 | Model predictions with and without gain shifting. 200 Hz time resolution. | N/A |
| **S2**<br>Stimulation<br>Suppression | 440<br>560<br>560 | 1<br>1<br>140 | 6.67-9.49<br>6.86-9.68<br>9.68 | Acquisition: 4 kHz low-pass filter, 10 kHz sampling. Additional low-pass filter: 10 Hz for persistent responses, 2-10 Hz for transient responses. | Persistent response: 10-s window 130-140 s after light stimulation. Transient response: average current in a 100-ms window centered on the response peak. Baseline subtracted (10-s window prior to the first light stimulus). |
| **S3A**<br>Induction<br>Suppression | 440<br>560 | 30<br>30 | 9.54<br>9.49 | Acquisition: 4 kHz low-pass filter, 10 kHz sampling. | N/A |
| **S3B**<br>Induction<br>Suppression | 436<br>577 | 0.2 or 10<br>10 | 10.09<br>10.97 | Acquisition: 10 kHz low-pass filter, 50 kHz sampling. | Transient response: Largest subthreshold depolarization during or shortly after illumination. Inflection: Most hyperpolarized voltage after the cessation of illumination. Persistent response: 5-s window immediately prior to the suppression pulse. Suppression: 5-s window $\geq 60$ s after the end of the suppression pulse. Baseline: 5-s window immediately prior to the first light stimulus. |
| **S8**<br>Stimulation<br>Suppression | Oc.-Filt. Xe<br>Xe<br>560 | 1<br>1<br>140 | 7.47-10.27<br>7.43-10.35<br>9.68 | Acquisition: 4 kHz low-pass filter, 10 kHz sampling. Additional low-pass filter: 10 Hz for persistent responses, 2-10 Hz for transient responses. | Persistent response: 10-s window 130-140 s after light stimulation. Transient response: average current in a 100-ms window centered on the response peak. Baseline subtracted (10-s window prior to the first light stimulus). |

**Table S2.**
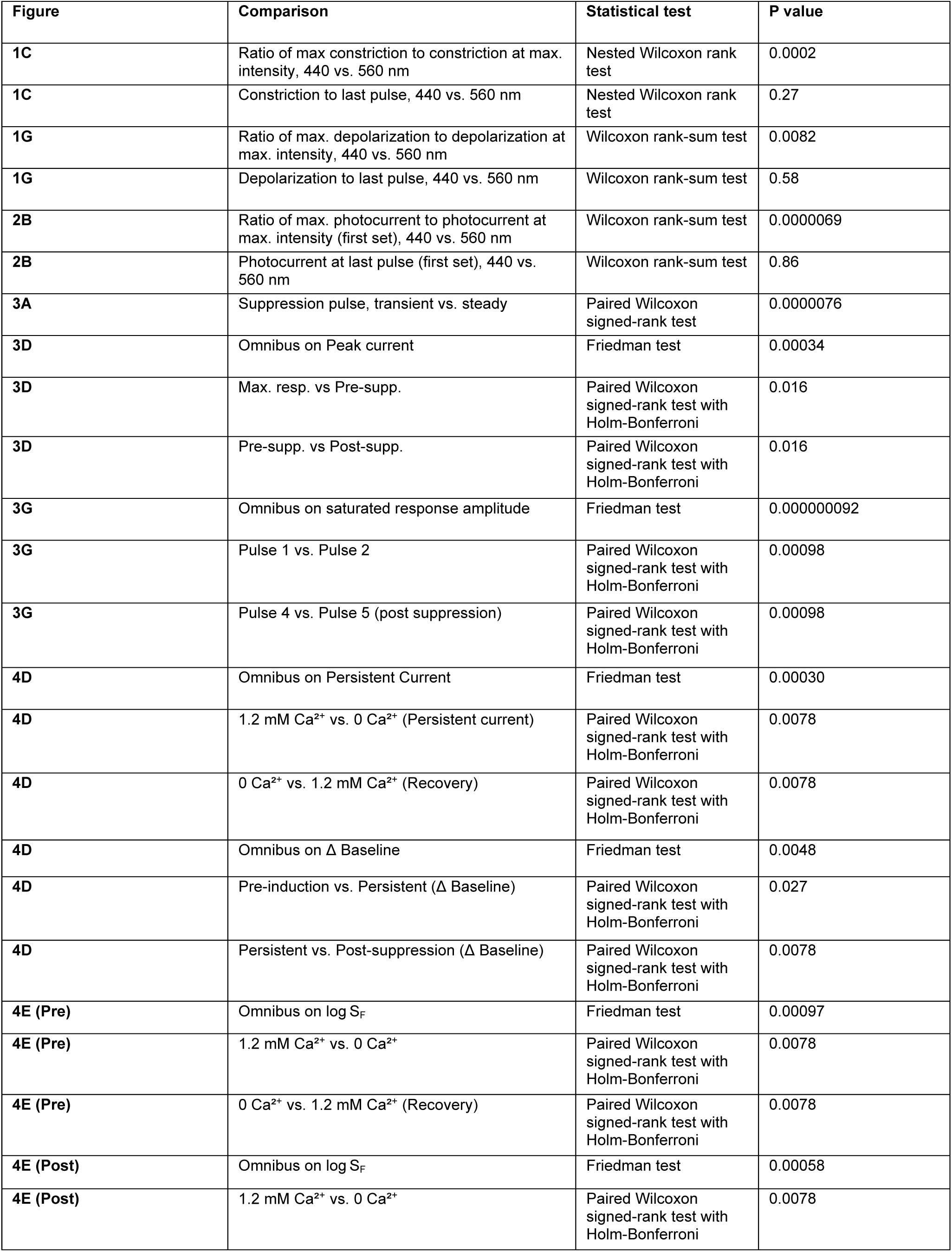
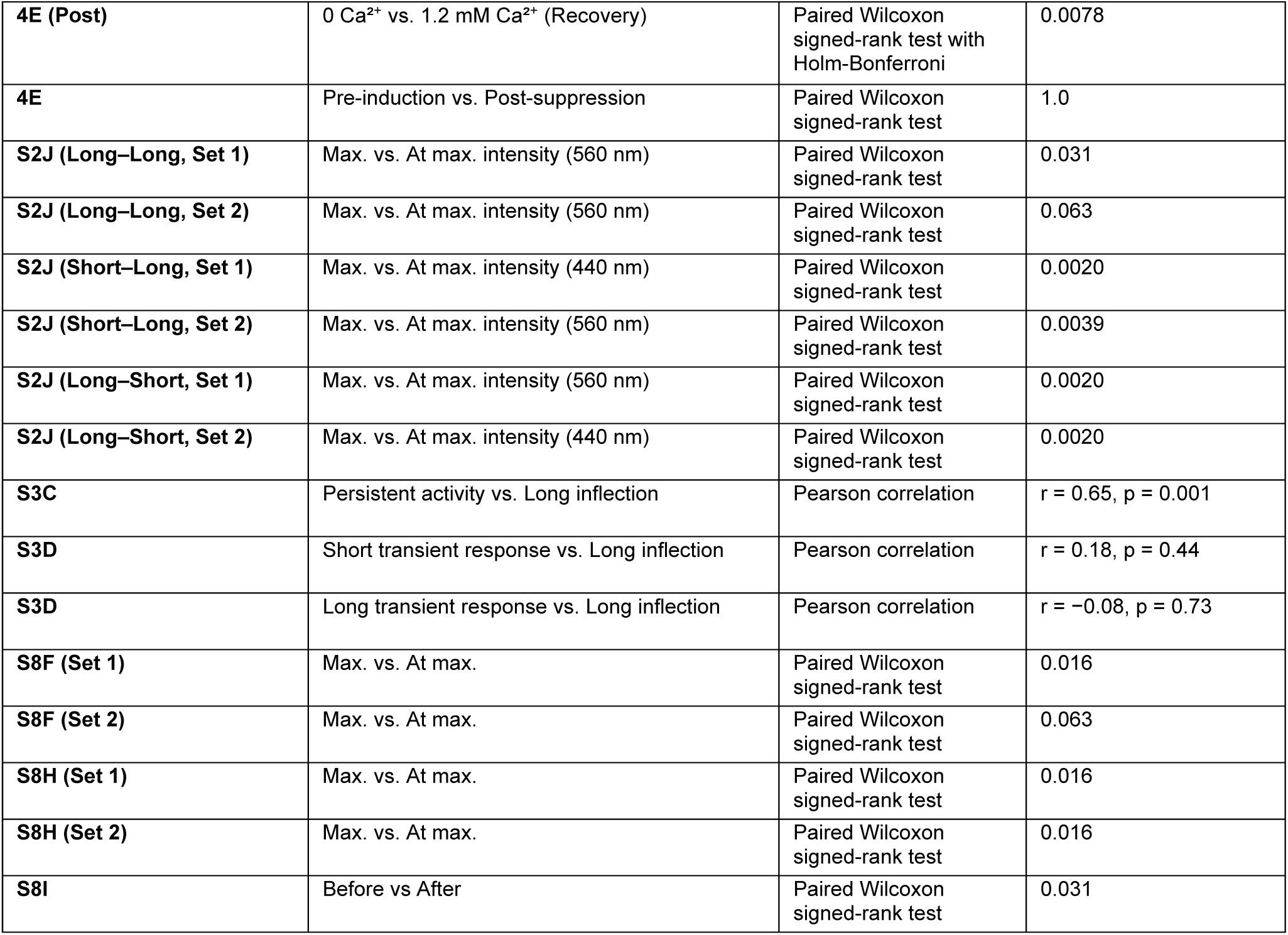
Statistical details. Summary of all statistical analyses associated with Figures 1-4, S2, S3, and S8. For paired comparisons, two-tailed Wilcoxon signed-rank tests were used to assess within-cell differences across experimental conditions. For correlation analyses, Pearson correlation coefficients (r) and corresponding p values are reported.

